# Structural basis of RNA-induced autoregulation of the DExH-type RNA helicase maleless

**DOI:** 10.1101/2022.11.11.516098

**Authors:** Pravin Kumar Ankush Jagtap, Marisa Müller, Anna E. Kiss, Andreas W. Thomae, Karine Lapouge, Martin Beck, Peter B. Becker, Janosch Hennig

## Abstract

Unwinding RNA secondary structures by RNA helicases is essential for RNA metabolism. How the basic unwinding reaction of DExH-type helicases is regulated by their accessory domains is unresolved. Here, we combine structural and functional analyses to address this challenge for the prototypic DExH RNA helicase maleless (MLE) from *Drosophila*.

We captured the helicase cycle of MLE with multiple structural snapshots. We discovered that initially, dsRBD2 flexibly samples substrate dsRNA and aligns it with the open helicase tunnel. Subsequently, dsRBD2 releases RNA and associates with the helicase core, leading to closure of the tunnel around ssRNA. Structure-based MLE mutations confirm the functional relevance of the structural model in cells. We propose a molecular model in which the dsRBD2 domain of MLE orchestrates large structural transitions that depend on substrate RNA but are independent of ATP.

Our findings reveal the fundamental mechanics of dsRNA unwinding by DExH helicases with high general relevance for dosage compensation and specific implications for MLE’s human orthologue DHX9/RHA mechanisms in disease.

## Introduction

RNA helicases are essential enzymes for remodelling RNA during a wide range of cellular processes including but not limited to pre-mRNA splicing, rRNA processing, translation, RNA transport, surveillance, and eventual degradation of the RNA (Bleichert and Baserga, 2007; Jankowsky, 2011; Jarmoskaite and Russell, 2014). These RNA-dependent ATPases use their helicase activity to interconvert RNA secondary structures (Iggo and Lane, 1989; Lorsch and Herschlag, 1998). The binding of substrates, ATP and RNA, triggers structural transitions to a closed “on” state, which is reversed to an open “off” state upon hydrolysis of the ATP and concomitant release of the RNA. Given their crucial role in various aspects of gene expression, it is not surprising that helicase activity is tightly regulated.

Members of the eukaryotic DExH/DEAH helicase family are multidomain proteins. Their common helicase module is formed by RecA1 and RecA2 domains and is frequently flanked by N- or C-terminal auxiliary domains (Cordin et al., 2012; He et al., 2010; Prabu et al., 2015; Schmitt et al., 2018; Schutz et al., 2010; Tauchert et al., 2017; Wagner et al., 1998; Walbott et al., 2010). These auxiliary domains commonly engage in specific or non-specific RNA interactions that contribute to substrate selectivity in the biological context (Ozgur et al., 2015; Rudolph and Klostermeier, 2015). Conditional conformational changes of auxiliary domains may install positive or negative autoregulation of the helicase activity (Absmeier et al., 2017; Absmeier et al., 2015a; Absmeier et al., 2015b; Chakrabarti et al., 2011; Collins et al., 2009; Floor et al., 2016; Gowravaram et al., 2018; Santos et al., 2012). Although genetic and biochemical studies of these helicases over the past decades, along with the recent structural studies in the context of spliceosomes, have significantly advanced our understanding, detailed molecular insights of how eukaryotic DExH/DEAH RNA helicases recognize double-stranded (ds) RNA substrates and the functional interplay between the auxiliary domains and the helicase module to bind and unwind the dsRNA have not been provided.

MLE is an RNA helicase of the DExH subfamily (Kuroda et al., 1991). It is well studied in the context of dosage compensation in *Drosophila melanogaster,* where a male-specific-lethal (MSL) dosage compensation complex (DCC) selectively targets and activates X-chromosomal genes to enhance transcription approximately two-fold. The MSL complex consists of the male-specific signature subunit MSL2, the scaffold protein MSL1, the epigenetic reader MSL3, the histone acetyltransferase MOF (‘males-absent on the first’), the RNA helicase *maleless* (MLE) and one of two long non-coding RNAs roX1 or roX2 (‘RNA on the X’) (Lucchesi and Kuroda, 2015; Samata and Akhtar, 2018). In the absence of roX RNA, male flies cannot sustain dosage compensation, a lethal condition (Deng and Meller, 2006). During the process of dosage compensation, MLE, putatively assisted by Unr (‘upstream of N-Ras’), binds conserved roX-box motifs and remodels roX RNAs to an alternative conformation, which is then incorporated into the MSL complex (Figure 1A) (Ilik et al., 2017; Ilik et al., 2013; Maenner et al., 2013; Militti et al., 2014; Müller et al., 2020). This RNA remodelling by MLE is essential for localization of the MSL complex to the X territory of the nucleus (Ilik *et al*., 2013; Maenner *et al*., 2013). Besides dosage compensation, MLE is also involved in RNA editing and siRNA processing (Cugusi et al., 2016; Reenan et al., 2000).

**Figure 1:**
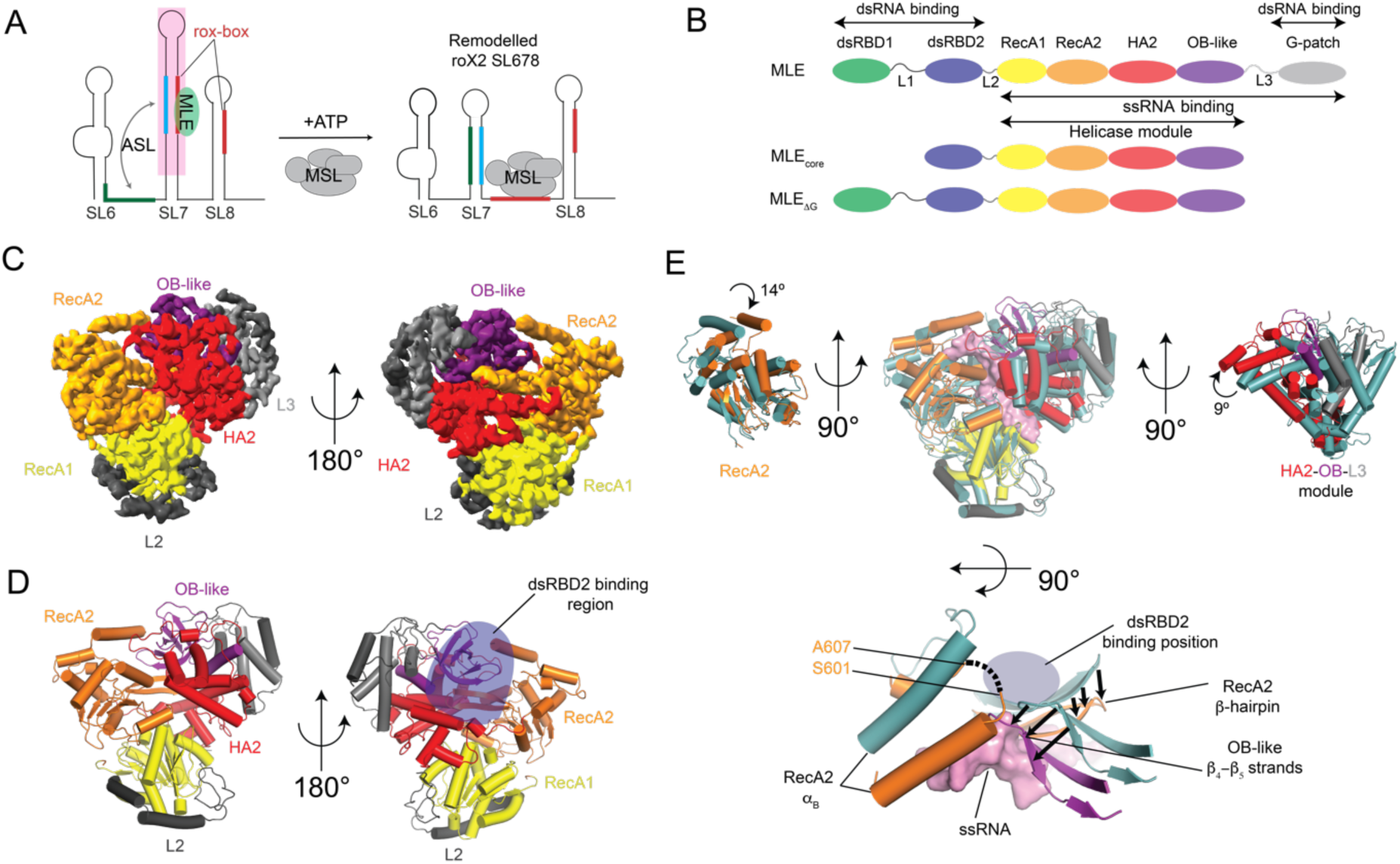
Cryo-EM structure of MLE_ΔG_ in the apo-state. (A) Schematic of the secondary structure of the biologically relevant three stem-loops in roX2 RNA located at the 3’ end. Upon remodelling of roX2 RNA by MLE, the MSL complex subunit MSL2 binds to the roX-box region. The region in roX2, of which dsRNA fragments were designed for cryo-EM studies with MLE is shaded in pink. (B) Domain organization of MLE from *D. melanogaster* and the constructs used for structural studies. The ss and dsRNA binding regions in MLE are indicated. (C) Cryo-EM density and (D) structure of MLE_ΔG_^apo^. The expected binding position of dsRBD2 on the MLE helicase module, as expected from the MLE_core_+U10 crystal structure, is marked with a blue oval. (E) Structural superposition of MLE_ΔG_^apo^ cryo-EM (with respective domain colours) and MLE_core_+U10 (light teal) crystal structures. Helices are shown as cylinders for simplicity. The structures were superposed with respect to the RecA1 domain. The rotations in RecA2 domain and HA2-OB-L3 module are shown. Black arrows in the top view indicate the inward movement of the RecA2 β-hairpin and the OB-like β_4_-β_5_ strands, which occludes the ssRNA binding tunnel in the MLE_ΔG_^apo^ structure. The missing loop between the RecA2 α_B_ and α_C_ is shown with dotted line in the MLE_ΔG_^apo^ structure.

MLE has a modular domain architecture consisting of two N-terminal auxiliary double-stranded RNA binding domains (dsRBDs), followed by the helicase domains (RecA1, RecA2), HA2 and OB-like domains and a C-terminal glycine-rich region (Figure 1B) (Prabu *et al*., 2015). This modular architecture segregates ss- and dsRNA binding properties between different parts of MLE. The dsRNA binding is mediated by the two N-terminal dsRBDs, while ssRNA is bound by the helicase module consisting of both RecA domains, the HA2 domain and the OB-like fold. The unstructured G-patch region binds to both ds- and ssRNA (Izzo et al., 2008). The sequence region encompassing dsRBD2 to the L3 linker forms a compact helicase core (MLE_core_) where the RecA, HA2, OB-like fold domains and the structured L3 linker form an RNA binding tunnel through which the ssRNA traverses during helicase translocation and thus form the helicase module of MLE (Prabu *et al*., 2015). Amongst the two dsRBDs, dsRBD2 was shown to be indispensable for helicase activity and localization to the X chromosome (Izzo *et al*., 2008).

The structures of the two tandem dsRBDs, free and upon dsRNA binding and the MLE_core_ in complex with a single-stranded uridyl-homopolymer RNA and ATP transition state analogue ADP:AlF_4_ have been determined previously (Jagtap et al., 2019; Lv et al., 2019; Prabu *et al*., 2015). However, these studies did not reveal the nature of RNA recognition by the dsRBDs in the context of the helicase and other auxiliary domains. Also, the structural rearrangements that characterize the productive RNA interactions of the apoenzyme, including binding of roX stem-loop (SL) structures, the funnelling of unwound ssRNA through the helicase and the role of auxiliary domains during dsRNA recognition and unwinding remain unknown. Here we address these questions using structural, biochemical, biophysical and functional studies supported by a series of cryo-EM structures of MLE at different states of its helicase cycle.

## Results

### MLE ssRNA binding tunnel is inaccessible in apo and nucleotide-bound states

A previously published crystal structure of MLE_core_ in complex with single-stranded RNA of 10 uridines (U10) and the ATP transition state analogue ADP:AlF_4_ (MLE_core_+U10) was interpreted as a closed state of the enzyme after unwinding the dsRNA substrate but still bound to ssRNA (Prabu *et al*., 2015). To determine the structural changes in MLE leading up to the substrate binding, we determined the cryo-EM structure of MLE_ΔG_ in the apo-state to a resolution of 3.8 Å (MLE_ΔG_^apo^)(Figure 1C-D, S1A-B, Table 1). Overall, the global architecture of MLE_ΔG_^apo^ in the cryo-EM structure resembles the MLE_core_+U10 crystal structure (Prabu *et al*., 2015). The helicase module adopts a trefoil arrangement formed by RecA1, RecA2 domains and the HA2-OB-L3 module. However, dsRBD2, which stably interacts with the RecA2 domain and the HA2-OB-L3 module in the MLE_core_+U10 crystal structure is not visible in the MLE_ΔG_^apo^ cryo-EM density map suggesting these two domains are flexible in the apo-state of the helicase.

A detailed comparison of the MLE_ΔG_^apo^ and MLE_core_+U10 structures revealed that ssRNA cannot be accommodated in the entry tunnel as it would clash with residues from the RecA2-α_B_ and α_B_-α_C_ loop in the MLE_ΔG_^apo^ structure (Figure 1E). The tunnel is further occluded by the loop between the HA2 α_6_-α_7_ helices and a β-hairpin formed by β_4_-β_5_ strands from the OB-like domain (Figure S2A). The HA2 loop residues extend into the entrance of the RNA tunnel and occupy positions corresponding to the 4^th^ and 5^th^ nucleotide of the RNA. Similarly, residues at the tip of the OB-like β_4_-β_5_ hairpin occupy positions corresponding to the 3^rd^ nucleotide in the MLE_core_+U10 structure (Figure S2A). In addition, RecA2 and HA2-OB-L3 modules show moderate rotation relative to the RecA1 domain upon binding of ssRNA. The RecA2 domain and the HA2-OB-L3 module rotate 14° and 9°, respectively, away from the RecA1 domain and towards the ssRNA binding tunnel (Movie S1).

As dsRBD2 is not visible in the MLE_ΔG_^apo^ structure, the region of the helicase module occupied by dsRBD2 at the back side of the RNA tunnel in the MLE_core_+U10 structure is solvent exposed. Residues 602-606 from the RecA2 domain, which are present in the distinctive insertion between α_B_ and α_C_ and interact with dsRBD2 in the MLE_core_+U10 structure are not visible and presumably flexible. The tip of the prominent RecA2 β-hairpin, present between β_4_ strand and α_4_ helix is pushed into the path of RNA in the tunnel. In the MLE_core_+U10 structure, this RecA2 β-hairpin forms interdomain protein-protein interactions with the dsRBD2 domain, which pulls the β-hairpin away from the RNA path (Figure 1E).

As the conformation of the RNA binding tunnel in MLE_ΔG_^apo^ would not support ssRNA binding, we wondered whether binding of ATP by the helicase module will promote structural rearrangements required for ssRNA binding in the tunnel. We thus determined the cryo-EM structure of MLE_ΔG_ in complex with ADP:AlF_4_ (MLE_ΔG_). We observed clear density for ADP:AlF_4_ in the nucleotide-binding pocket (Figure S2B). Interestingly, this structure showed only minor conformational changes compared to the MLE_ΔG_ structure with RecA2 moving by 3° towards RecA1, and HA2-OB-L3 moving approximately 4° away from RecA1 and towards RecA2 (Figure S2C). In addition, the distances between the centre-of-mass for the RecA1, RecA2 and HA2-OB-L3 module changed within ∼1 Å suggesting binding of ATP does not lead to an opening of the helicase module to allow ssRNA binding (Figure S2D). On the contrary, the tip of the β_4_-β_5_ hairpin from the OB-like domain moves even further into the RNA binding tunnel, forming a physical barrier to prevent ssRNA binding and promoting the closed conformation of MLE (Figure S2C).

An increase in the centre-of-mass distances between RecA1 and RecA2 domains have been correlated with the opening of the helicase module competent to bind ssRNAs in the RNA helicases (Hamann et al., 2019). In MLE_ΔG_^apo^, the RecA1-RecA2 centre-of-mass distance is 32.5 Å and does not change significantly upon binding of ADP:AlF_4_ as well as U10 ssRNA (Figure S2D). However, the centre-of-mass distance between the RecA2 domain and HA2-OB-L3 module increases by 5.3 Å upon binding of U10 RNA compared to the MLE_ΔG_^apo^ structure as opposed to a minor ∼1 Å in the MLE_ΔG_ structure. Clearly, in the case of MLE, the centre-of-mass distances between RecA2 domain and the HA2-OB-L3 module serves as a better metric to determine the opening of the ssRNA binding tunnel. Accordingly, the helicase module is in a closed conformation with an inaccessible RNA binding tunnel, independent of nucleotide binding. This contrasts other DExH RNA helicases including spliceosome-associated DEAH helicases Prp43 and Prp22, which show conserved residues involved in RNA and ATP-binding and yet reside in an open conformation competent for ssRNA binding with widely spaced RecA1 and RecA2 domains in the absence of adenosine nucleotides. In Prp22, the presence of adenosine nucleotides stabilize the closed conformation of the helicase module regardless of the presence of RNA (Caruthers et al., 2000; Hamann *et al*., 2019; Montpetit et al., 2011; Schmitt *et al*., 2018; Tauchert et al., 2016; Tauchert *et al*., 2017; Theissen et al., 2008).

### Structure of MLE in complex with dsRNA reveals opening of the helicase module upon dsRNA binding

The structures discussed above only consider MLE conformations either in the absence of RNA or bound to ssRNA. However, MLE is thought to initially recognize dsRNA in form of the 3’ SL7/8 of roX (Müller *et al*., 2020). While it is clear that the dsRBD1,2 domains of MLE are involved in this recognition, their structure in complex with dsRNA only involved isolated domains (Izzo *et al*., 2008; Jagtap *et al*., 2019; Lv *et al*., 2019; Prabu *et al*., 2015). dsRNA was predicted to bind to the MLE helicase module in a perpendicular position relative to the ssRNA involving structural changes of RecA2-α_B_ and α_B_-α_C_ loop to expose the RNA entry tunnel (Lv *et al*., 2019; Prabu *et al*., 2015).

To understand the mode of dsRNA binding to MLE, we attempted to solve the cryo-EM structure of MLE_ΔG_ in complex with roX2 stem-loop 678 (SL678, Figure 1A). However, samples were heterogeneous as observed in EMSA experiments and the complex dissociated on cryo-EM grids. The heterogeneity primarily arose from the formation of multimers due to several MLE molecules binding to the long RNA (Figure S3A). To circumvent this, we designed a dsRNA derived from the stem-loop 7 of roX2 with a 3’ single-stranded extension consisting of the UxUUU ssRNA motif shown to bind MLE in the RNA tunnel (Prabu *et al*., 2015). This RNA (called ‘SL7’ hereafter) consisted of a stem of 19 base pairs for dsRBD1,2 binding and a 14 nucleotide 3’ extension of U-rich ssRNA intended to fit into the helicase module. MLE_ΔG_ bound to this RNA with an affinity of 16 nM, and the complex of MLE_ΔG_ with SL7 RNA showed the formation of a single species in EMSA experiments (Figure S3B-E).

We obtained a cryo-EM structure of MLE_ΔG_ in complex with SL7 RNA and ADP:AlF_4_ (MLE_ΔG_+SL7) at a resolution of 4 Å (Figure 2A-B, Figure S1, Table 1). Remarkably, the dsRBD2 domain bound to dsRNA is now visible in the EM density. The dsRBD2 aligns the dsRNA with the ssRNA binding tunnel as opposed to a perpendicular conformation predicted earlier (Figure 2A-B). Residues from the loop connecting β_1_-β_2_ strands of dsRBD2 interact with the RNA backbone from the 2^nd^ minor groove (counting from the proximal base relative to the helicase module). The α_1_ helix of dsRBD2 binds to the first minor groove, and the N-terminal residues of α_2_-helix bind to the next major groove from the base of the stem. These canonical dsRBD-dsRNA interactions agree with previous structural data (Jagtap *et al*., 2019; Lv *et al*., 2019; Masliah et al., 2013). N-terminal residues at the L2 linker causes structural constraints and defines the range within which dsRNA can be contacted to the 1st register of the minor-major-minor groove (Figure 2B).

**Figure 2:**
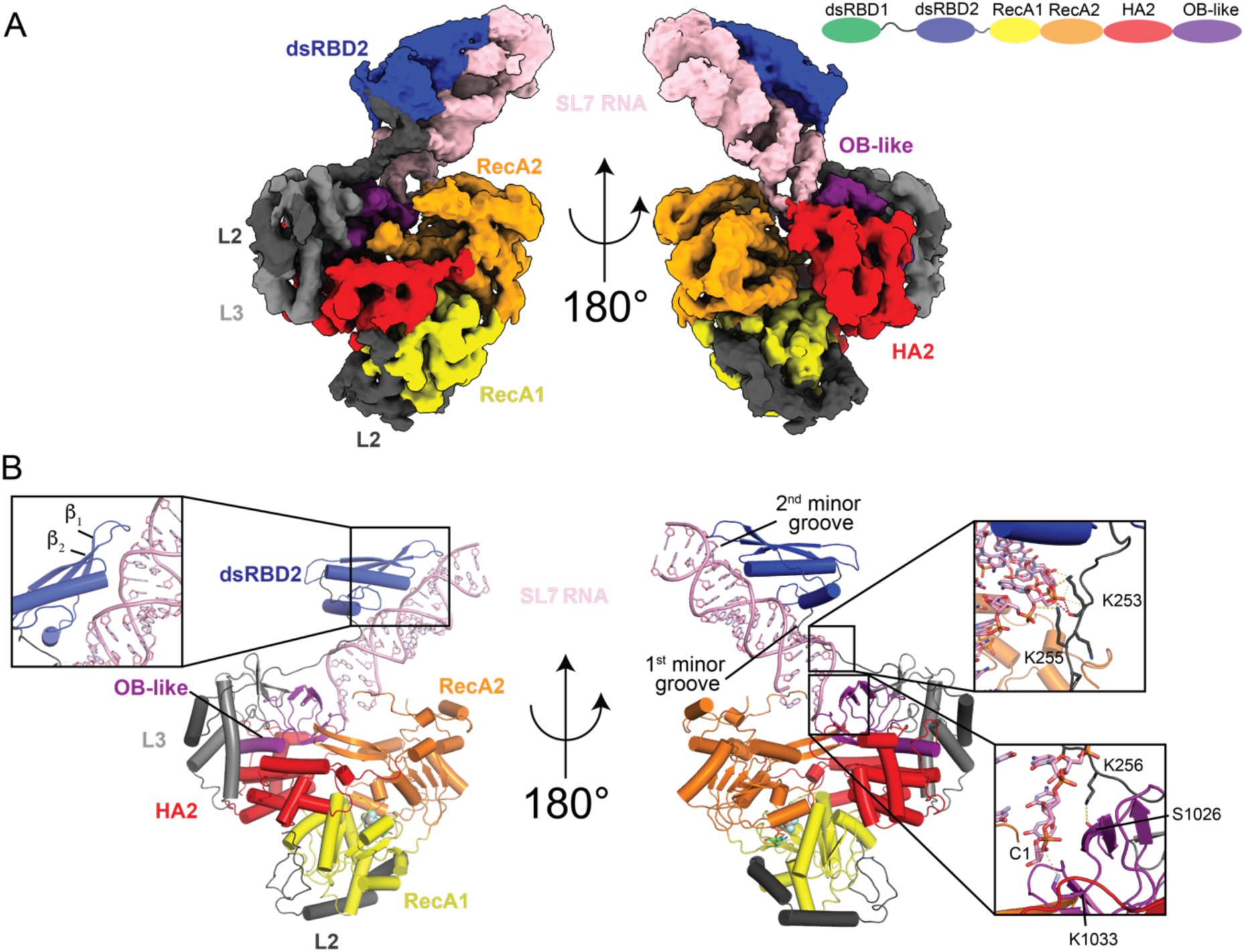
Structure of MLE_ΔG_ in complex with dsRNA. (A) Cryo-EM density map of MLE_ΔG_+SL7 with respective domain colours. The dsRNA is shown in pink. (B) Structure of MLE_ΔG_+SL7. Zoomed-in panels show key interactions between the RNA and different domains of MLE. The two minor groves in the dsRNA are marked. Yellow and red dashed lines show polar and electrostatic contacts respectively.

Several features of dsRNA recognition by MLE are noteworthy. dsRBD2 along with the N-terminal 26 residues of the L2 linker provide a continuous patch of positive electrostatic surface potential which interacts with the negatively charged dsRNA backbone. This positively charged surface continues into the RNA entry tunnel lined with residues from the OB-like and HA2 domains (Figure S3G). Two of the four L2 linker lysines (K253 and K255) interacting with the phosphate backbone of dsRNA guide the 3’-end of the dsRNA to the entry tunnel of the helicase module. The side-chain amine of K256 from this linker further forms a hydrogen bond with the backbone carbonyl of S1026 in the OB-like domain, thereby restricting the mobility of the linker and anchoring it to the helicase module (Figure 2B). The lysines 254-256 are flexible in the MLE_core_+U10 crystal structure and in the isolated dsRBD1,2 domains, suggesting that these residues only become ordered in the presence of dsRNA (Prabu *et al*., 2015). This mode of dsRNA recognition and presumed funnelling of unwound RNA into the tunnel appears to be generally relevant, since K253-255 are highly conserved amongst different species (Figure S3H).

Our novel structure reveals large conformational changes of MLE upon dsRNA binding. To accommodate dsRNA at the entry of the ssRNA binding tunnel, the RecA2 domain and the HA2-OB-L3 module rotate by 14° and 19°, respectively, away from the RNA entry tunnel compared to the MLE_ΔG_ structure (Figure 3A, S4A, Movie S2). Near the RNA entry tunnel, these rotations cause the retraction of the HA2 α_6_-α_7_ loop and α_10_ helix and the RecA2 α_2_/α_3_ helices, allowing the 3’-end of the dsRNA to approach the entry tunnel. Furthermore, the RecA2 α_B_ helix is not visible in the EM-density suggesting it is flexible in the dsRNA bound structure, thereby allowing to accommodate the 5’-end of the dsRNA (Figure 3B).

**Figure 3:**
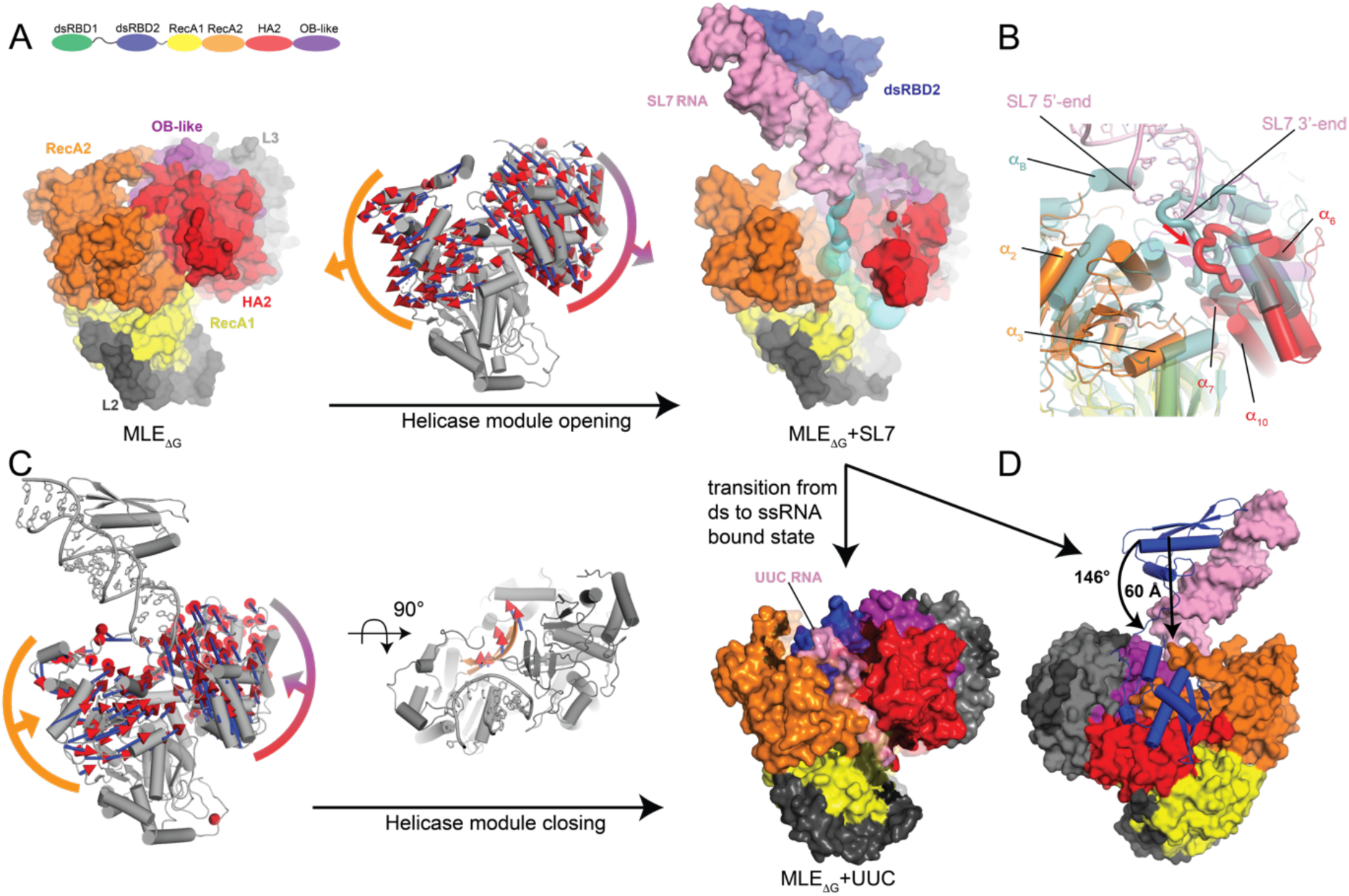
Opening and closing of MLE helicase module upon binding to ds and ssRNA. (A) The MLE_ΔG_ and MLE_ΔG_+SL7 structures are shown as surface representations. Movement of RecA2 domain and HA2-OB-L3 module upon dsRNA binding and opening of the helicase module is shown with C_α_ vectors from blue to red on the cartoon representation of MLE_ΔG_ structure (in grey, with RecA2 and HA2-OB-L3 module shown with orange and red-purple-grey arcs). The formed RNA binding tunnel in MLE_ΔG_+SL7 structures is shown in cyan. The neighbouring region around the RNA binding tunnel is shown with a transparent surface for clear visibility of the tunnel. (B) Zoom-in of the RNA binding tunnel showing displacement of structural elements from RecA2 and HA2 domain upon binding of dsRNA to MLE_ΔG_ (with respective domain colours) compared to MLE_ΔG_ (cyan) structure. The loop between α_6_-α_7_ helices from OB-like domain is shown with thicker radius for clarity with its motion shown by red arrow. (C) Closing of the helicase module upon transition from SL7 dsRNA to UUC ssRNA bound state. The movement of RecA2 domain and HA2-OB-L3 module upon transition from ds to ssRNA bound state is shown with C_α_ vectors from blue to red on the cartoon representation of MLE_ΔG_+SL7 structure (in grey). The 90° top view depicts the movement of RecA2 β-hairpin (orange) out of the ssRNA binding tunnel. The UUC ssRNA bound state of MLE_ΔG_ is shown as surface representation with ssRNA in pink and neighbouring region around ssRNA in transparent for clarity. (D) Flipping of the dsRBD2 onto the helicase module is shown on the MLE_ΔG_+SL7 structure upon transition from ssRNA to dsRNA bound state.

### Conformational rearrangement of dsRBD2 drives binding of ssRNA into the helicase module

Surprisingly, we did not observe the density of ssRNA in the MLE_ΔG_+SL7 structure, although the U-rich single-stranded extension at the 3’-end of the dsRNA was designed to be accommodated in the tunnel. To assure that the missing ssRNA in our structure was not due to the inability of the helicase module to bind RNA sequences other than poly-U, we determined the cryo-EM structures of MLE_ΔG_ in complex with ADP:AlF_4_ and either U10-mer ssRNA (MLE_ΔG_+U10 structure) or a UUC-rich ssRNA (MLE_ΔG_+UUC structure) derived from the ss extension of SL7 RNA (CCUCUUUCUUUC, Figure S1, S3I-L). Both structures superpose well with the MLE_core_+U10 crystal structures with a root mean square deviation of 0.7 Å and 1.4 Å, respectively. We could observe density for the single-stranded U10 and UUC RNAs in the RNA binding tunnel of the helicase module. Contrary to the MLE_ΔG_^apo^ structure (Figure 1), dsRBD2 is now visible and occupies the same position on the RecA2 β-hairpin and HA2-OB-L3 surface of the helicase module as in the MLE_core_+U10 structure. Furthermore, dsRBD1 and the linker connecting dsRBD1,2 are not visible, confirming that this region remains flexible in the presence of ssRNA.

In summary, MLE_ΔG_ is capable of binding to UUC ssRNA as evident from the cryo-EM structure, but the ssRNA moiety of SL7 RNA is not bound. We further analysed the MLE_ΔG_+SL7 structure to find an explanation for this conundrum. Although the RNA binding tunnel is accessible to ssRNA, Tyr752 of the RecA2 β-hairpin and the OB-like β-hairpin formed by β_4_-β_5_ strands clash with the putative binding sites of nucleotides 3 to 7 (Figure S4B). Comparison of MLE_ΔG_+SL7 and MLE_ΔG_+UUC structures revealed that for ssRNA insertion into the helicase module and transition from dsRNA to ssRNA bound state, the RecA2 domain and HA2-OB-L3 module must move inward towards the ssRNA binding tunnel accompanied by rotations of 16° and 17°, respectively (Figure 3C, S4C, Movie S3). These motions of RecA2 domain and HA2-OB-L3 module close the helicase module and lead to binding of ssRNA into the helicase module.

These domain motions are in turn due to an extensive conformational change of dsRBD2 (Figure 3D, S4C). To bind ssRNA, dsRBD2 flips back onto the helicase module with a displacement of 60 Å, while rotating by 146°. For simplicity, we call the conformation of dsRBD2 in MLE_ΔG_+SL7 structure “flipped-out” (‘out’ in construct names) and in complex with ssRNA as “flipped-in” (‘in’ in construct names) conformations. In the flipped-in conformation, dsRBD2 acts as a brace and holds together the RecA2 domain and the HA2-OB-L3 modules by forming an intricate network of hydrogen bonds and hydrophobic interactions (Figure S4D-E for structural details). These interactions help to pull the RecA2 β-hairpin out of the ssRNA path within the RNA tunnel (Figure 3C, S4E). All these structural changes caused by the interaction of the dsRBD2 with the helicase module stabilize the ssRNA bound closed state of MLE.

### The interactions of dsRBD1,2 with the helicase module are dynamic

Although the construct used for cryo-EM studies contained dsRBD1, we did not observe any density for it in any of our cryo-EM structures, suggesting that dsRBD1 remains flexible in these states. This supports earlier conclusions that dsRBD1 binds weakly to dsRNA and the linker connecting dsRBD1,2 remains flexible upon RNA binding (Izzo *et al*., 2008; Jagtap *et al*., 2019).To determine if there are any transient interactions between dsRBD1 and the helicase module which could not be captured in our cryo-EM structures, and to recapitulate the interaction of dsRBD2 with the helicase module, we monitored changes of the ^1^H-^15^N HSQC NMR spectrum of ^15^N-labelled dsRBD1,2 upon addition of an equimolar concentration of the unlabelled helicase module. To our surprise, amide peaks of dsRBD1 showed strong line broadening and consequently a decrease in signal intensity compared to minor line broadening in the linker and dsRBD2 (Figure 4A-B, S5A, Table S1). The line broadening is prominently located on the three β-strands and to some extent on the two α-helices for both dsRBD1 and dsRBD2 (Figure 4C). This suggests that dsRBD1 interacts stronger with the helicase module than dsRBD2, an unexpected finding given published data and our cryo-EM structures (Prabu *et al*., 2015).

**Figure 4:**
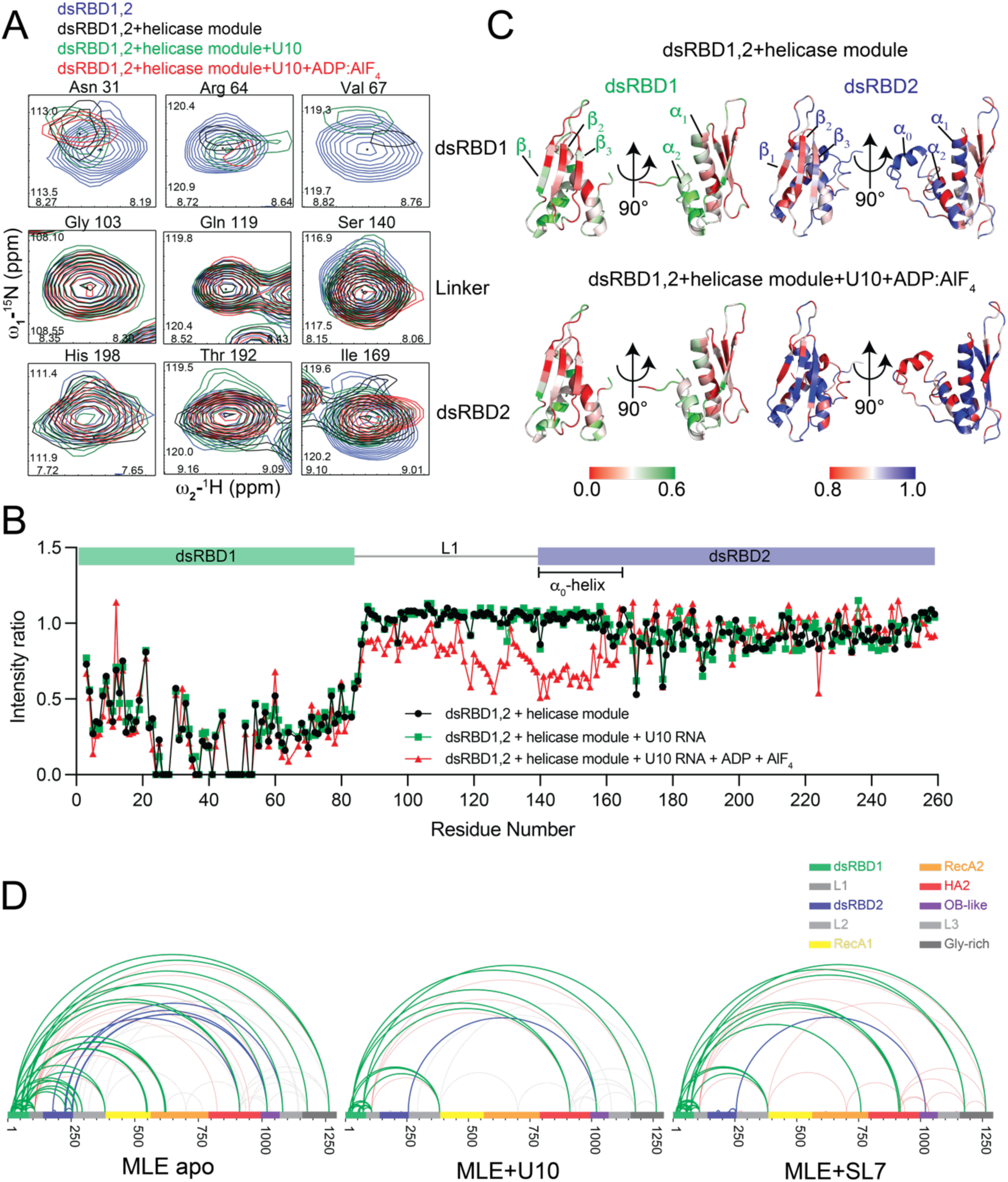
Interaction between MLE dsRBD1,2 and the helicase module. (A) Individual exemplary peaks from dsRBD1, linker and dsRBD2 region from the superposed ^1^H,^15^N HSQC NMR spectrum of dsRBD1,2 free and bound to equimolar amount of helicase module in the presence and absence of ss U10 RNA and ADP:AlF_4_. (B) Intensity ratios of dsRBD1,2 as quantified from spectra shown in Figure S5A. See also table S1 for average intensity decrease in the individual domains. (C) Mapping of line broadening and intensity decrease of NMR signals in dsRBD1,2 plotted on the NMR structures of dsRBD1 and dsRBD2 upon titration with equimolar ratio of MLE helicase module alone and in complex with ss U10 RNA and ADP-AlF_4_. (D) Interdomain crosslinks within MLE in the apo-state, in complex with U10 ssRNA or with SL7 dsRNA are presented as arches. Crosslinks originating from dsRBD1, linker and the dsRBD2 are shown in green, red, and blue for MLE, respectively.

To understand if the presence of RNA and nucleotides modulates the interaction between dsRBD1,2 and the helicase module, we performed NMR titration experiments in the presence of U10 ssRNA and ADP:AlF_4_. Although ssRNA itself did not have any significant effect on the interaction between dsRBD1,2 and helicase module, the presence of ssRNA and ADP:AlF_4_ led to an intensity decrease in linker residues and in the α_0_ helix of dsRBD2. The observed line broadening is consistent with our previous findings that this dynamic α_0_ helix does not bind RNA but interacts with the helicase module in the MLE crystal structure in complex with U10 and ADP:AlF_4_ (MLE_core_+U10) (Jagtap *et al*., 2019; Prabu *et al*., 2015). By contrast, line broadening of dsRBD1 residues is unaffected by the presence of ADP:AlF_4_ and ssRNA (Figure 4A-C, S5A, Table S1).

To resolve the apparent contradiction between NMR and cryo-EM data with regards to dsRBD – helicase interactions, we determined the proximity of amino acids in full-length MLE apoenzyme and in the presence of ssRNA and dsRNA by cross-linking mass spectrometry (CL-MS) (Figure 4D, Table S2). In the apo-state, 7 lysines from dsRBD1 and a single lysine from the linker region crosslink with 8 lysines from the helicase module and the G-patch. However, dsRBD2 lysine residues only crosslinked with three lysines, K936, K1020 and K1081, located in HA2, OB-like and L3 regions of the helicase module. Two of these lysines (K1020 and K1081) are within crosslinking distance of the previously known binding site of dsRBD2, as demonstrated by the cryo-EM structure of MLE_ΔG_ (Figure S5B). In the presence of U10 ssRNA or SL7 dsRNA, the dsRBD1-linker formed fewer crosslinks, but with similar distribution. On the other hand, K256 from dsRBD2 formed only one specific interdomain crosslink with K1020 of the OB-like domain. From our cryo-EM structures, this crosslink is possible if MLE binds to either ss or dsRNA. Therefore, these data are in complete agreement with our cryo-EM structures and suggest that dsRBD1-linker interacts non-specifically with the MLE core independent of the presence of RNA and adenosine nucleotides, whereas dsRBD2 has limited flexibility in the absence of RNA and assumes a specific conformation in the presence of RNA.

### dsRBD2 conformations modulate RNA binding and helicase activity in *cis*

So far, we showed that in the dsRBD2 ‘flipped-out’ conformation, MLE exists in an open conformation and binds dsRNA, and in the dsRBD2 ‘flipped-in’ conformation, MLE exists in a closed conformation and binds ssRNA. To validate these findings, we designed structure-based mutations predicted to lock MLE either in the dsRBD2 ‘flipped-in’ or ‘flipped-out’ conformation. To stabilize the ‘flipped-in’ conformation, thus forcing the closed conformation of MLE (called MLE_ΔG_^in^), we engineered a cysteine bridge by mutating Glu195 of dsRBD2 and Ser633 of RecA2 to cysteines. To force dsRBD2 in the flipped-out conformation we mutated 13 residues to alanine to disrupt all hydrogen bonds between dsRBD2 and the helicase module to detach dsRBD2 (called MLE_ΔG_^out^). Both mutants were properly folded in solution as determined by circular dichroism spectroscopy (Figure S6A). In the absence of the reducing agent TCEP MLE_ΔG_^in^, like unmutated MLE_ΔG_ was mildly aggregated, as seen from the size exclusion chromatography coupled with multi-angle light scattering (SEC-MALS) (Figure S6B). In addition, the melting temperature for both mutant proteins was within ±3 °C of the wild-type protein, further confirming that the stability of the two mutants does not change significantly (Figure S6C). The melting temperature of the MLE_ΔG_^in^ mutant increased by 2.5 °C suggesting the formation of a disulphide bond between Cys195-Cys633, thereby stabilising MLE in the dsRBD2 flipped-in conformation.

We tested both mutants for binding to either roX2 SL678 RNA or UUC ssRNA by fluorescence polarisation assays (Figure 5A, Table S3). MLE_ΔG_^WT^ and MLE_ΔG_^out^ bound to the SL678 with an affinity of 4.2 and 8.9 nM compared to MLE_ΔG_^in^, which showed a decreased dsRNA affinity by ∼20-fold (80 nM). In contrast, MLE_ΔG_^out^ showed a decreased ssRNA affinity (>2500 nM) compared to MLE_ΔG_^WT^ (302 nM). MLE_ΔG_^in^ bound to UUC RNA similarly to MLE_ΔG_^WT^ (280 nM). These experiments suggest that the dsRBD conformation modulates the preference of MLE for binding to ss or dsRNA.

**Figure 5:**
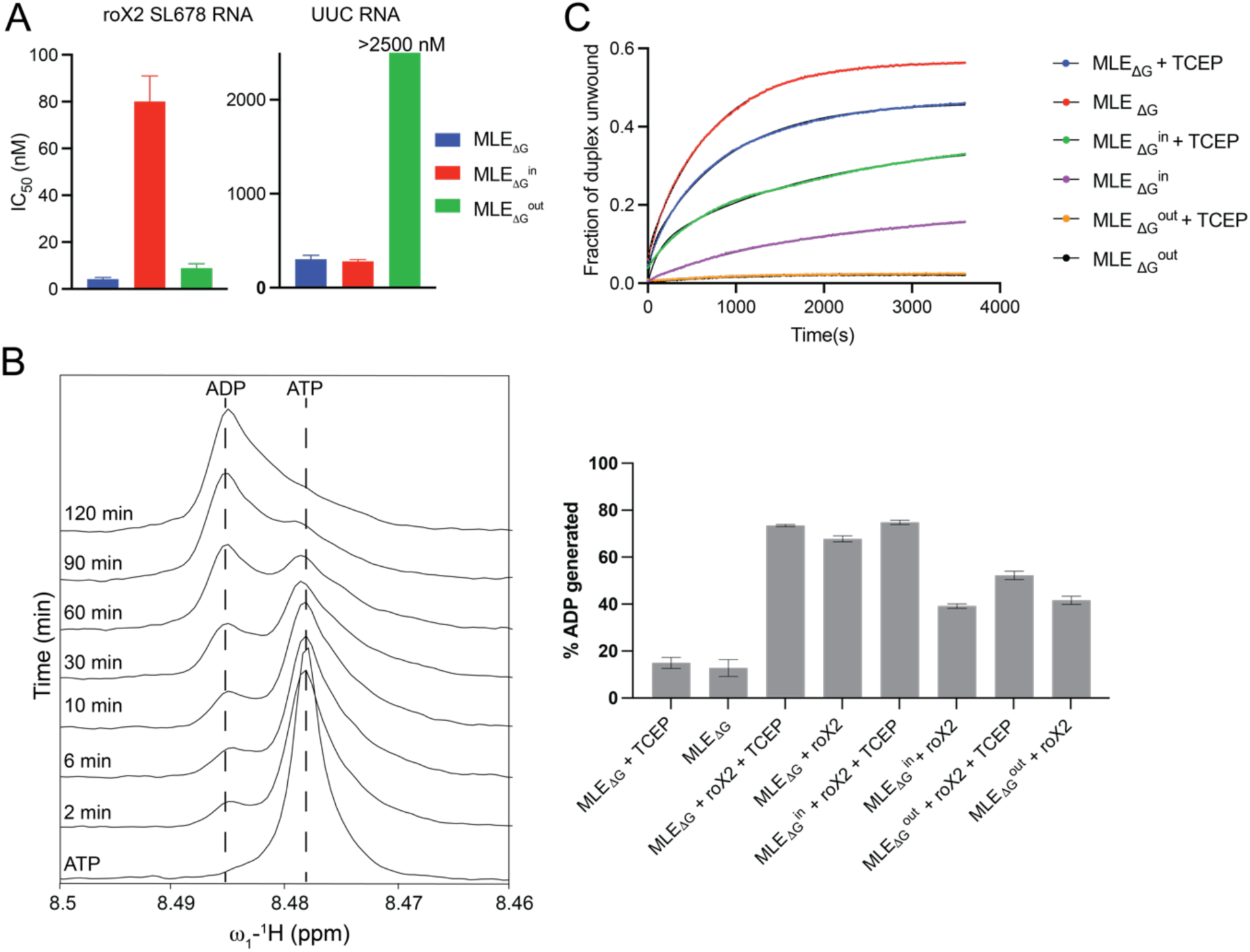
Biophysical and biochemical characterization of MLE mutants. (A) Affinity of MLE_ΔG_ wild type and mutants for roX2 SL678 and UUC RNA shown as half-maximal inhibitory concentration (IC_50_) as determined by fluorescence polarization assays. (B) NMR 1D ATPase assays of MLE_ΔG_ and its mutants. ^1^H NMR curves at exemplary time points (left) and the percentage of ADP generated after 120 min (right) are presented. (C) Helicase activity of MLE_ΔG_ and its mutants as a function of time. The curves were fit to a two-phase association. The timepoint at which ATP was added was adjusted to zero after recording baseline fluorescence for 4 mins. The assays were recorded in triplicates.

To assess the effect of these mutations on the helicase activity of MLE, we first confirmed that the MLE mutants retain their ATPase activity by measuring the rate of ATP to ADP conversion using 1D NMR experiments (Figure 5B). The ATPase activity of DExH helicases is stimulated by the presence of RNA (Kim et al., 1992; Schwer and Guthrie, 1991; Tanaka and Schwer, 2005; 2006). As expected, MLE_ΔG_^WT^ did not show significant ATPase activity in the absence of SL678 RNA, suggesting that MLE only hydrolyses ATP when bound to RNA. In the presence of SL678 RNA, MLE_ΔG_^in^ and MLE_ΔG_^out^ showed a slight decrease in their ATPase activity compared to MLE_ΔG_^WT^ (Figure 5B, Table S4). In the presence of TCEP, MLE_ΔG_^in^ regained full ability to hydrolyse ATP equivalent to MLE_ΔG_^WT^, suggesting the release of dsRBD2 from the helicase module due to TCEP reducing the disulphide bridge. Overall, these results indicate that these MLE mutants are able to bind and hydrolyse ATP.

Next, we assessed the real-time helicase activity of these mutants using a model duplex RNA derived from SL7 RNA used for cryo-EM and labelled with black hole quencher-1 (BHQ-1) at the 3’-end of the 1^st^ strand and 6-carboxyfluorescein (6-FAM) at the 5’-end of the 2^nd^ strand (Figure 5C, S3F). Upon annealing the two strands, the 6’-FAM fluorescence signal is quenched by close-by BHQ-1. An increase of fluorescence signal intensity upon addition of MLE reveals its helicase activity. In these experiments, the steady state RNA fraction unwound by MLE_ΔG_^in^ (18%) is 2.5-fold lower than the MLE_ΔG_^WT^ (46%) suggesting it unwinds dsRNA albeit with lower efficiency. Upon addition of TCEP to release the dsRBD2 from the helicase module, MLE_ΔG_^in^ regained its activity with only a minor 1.2-fold decrease in efficiency compared to MLE_ΔG_^WT^. However, the ability of MLE to unwind dsRNA was almost completely lost in the MLE_ΔG_^out^ mutant, with only ∼2% of the RNA unwound under steady state conditions (Table S5).

### Restricting dsRBD2 conformations affects X chromosome localisation, territory formation and RNA binding *in vivo*

Next, we assessed the relevance of dsRBD2 conformation for MLE’s *in vivo* function. MLE colocalizes with the MSL-DCC (DCC for short) and roX RNA on the male X chromosome, where they bind to MRE sequences at active genes. Faithful localisation to the X chromosome is revealed as staining of a coherent X chromosome territory by immunofluorescence microscopy. MLE binding to the territory depends on its interaction with roX2 (Prabu *et al*., 2015) and conversely, coherent territory staining depends on the presence of MLE and its ATPase and helicase activities (Gu et al., 2000). We generated S2 cell lines stably expressing GFP-tagged, full-length wildtype MLE (MLE_fl_^WT^) or dsRBD2 mutants MLE_fl_^in^ and MLE_fl_^out^, respectively. The *mle* transgenes had been rendered RNAi-resistant by synonymous codon adaption, allowing to monitor their function after depletion of endogenous MLE. RNAi directed against irrelevant glutathion-S-transferase (GST) sequences served as control (Figure S7A). The rescue capability of GFP-tagged transgenes was monitored by qualitative and quantitative immunofluorescence microscopy (Figure S7B). The cell lines were non-clonal and the GFP-negative cell pool served as internal negative control. As expected, depletion of MLE by RNAi impaired X chromosome territory formation, which also translated into an overall drop of MSL2 levels (panel S2 *mle* RNAi) (Figure 6A-B, S7C). Expression of MLE_fl_^WT^-eGFP restores MSL2-marked territories, the transgene localizes to X-chromosome territories and MSL2 nuclear levels are also increased relative to the GFP-negative cell pool (Figure 6A-B, S7C). A comparable phenotype was observed in MLE_fl_^in^-eGFP-expressing cells, which seems in contrast to the observed deficiencies of this mutant *in vitro* (Figure 5). However, due to the intracellular reducing environment the intended disulfide bonds in this mutant are most likely unstable *in vivo*, rendering MLE_fl_^in^-eGFP a wild-type like variant. This result also demonstrates that the cysteine substitutions *per se* do not alter the physiological function of the enzyme. In contrast, we observed a strong phenotype in MLE_fl_^out^-eGFP-expressing cells (Figure 6A-B, S7C). In control *gst* RNAi-treated cells endogenous MLE still localized to MSL2-stained X territories while MLE_fl_^out^-eGFP did not colocalize and exhibited a diffuse and speckled nuclear localization. In absence of endogenous MLE, the MLE_fl_^out^-eGFP mutant failed to rescue X chromosome territory formation and restore physiological MSL2 protein levels. The data confirm that the helicase-deficient enzyme does not function *in vivo*.

**Figure 6:**
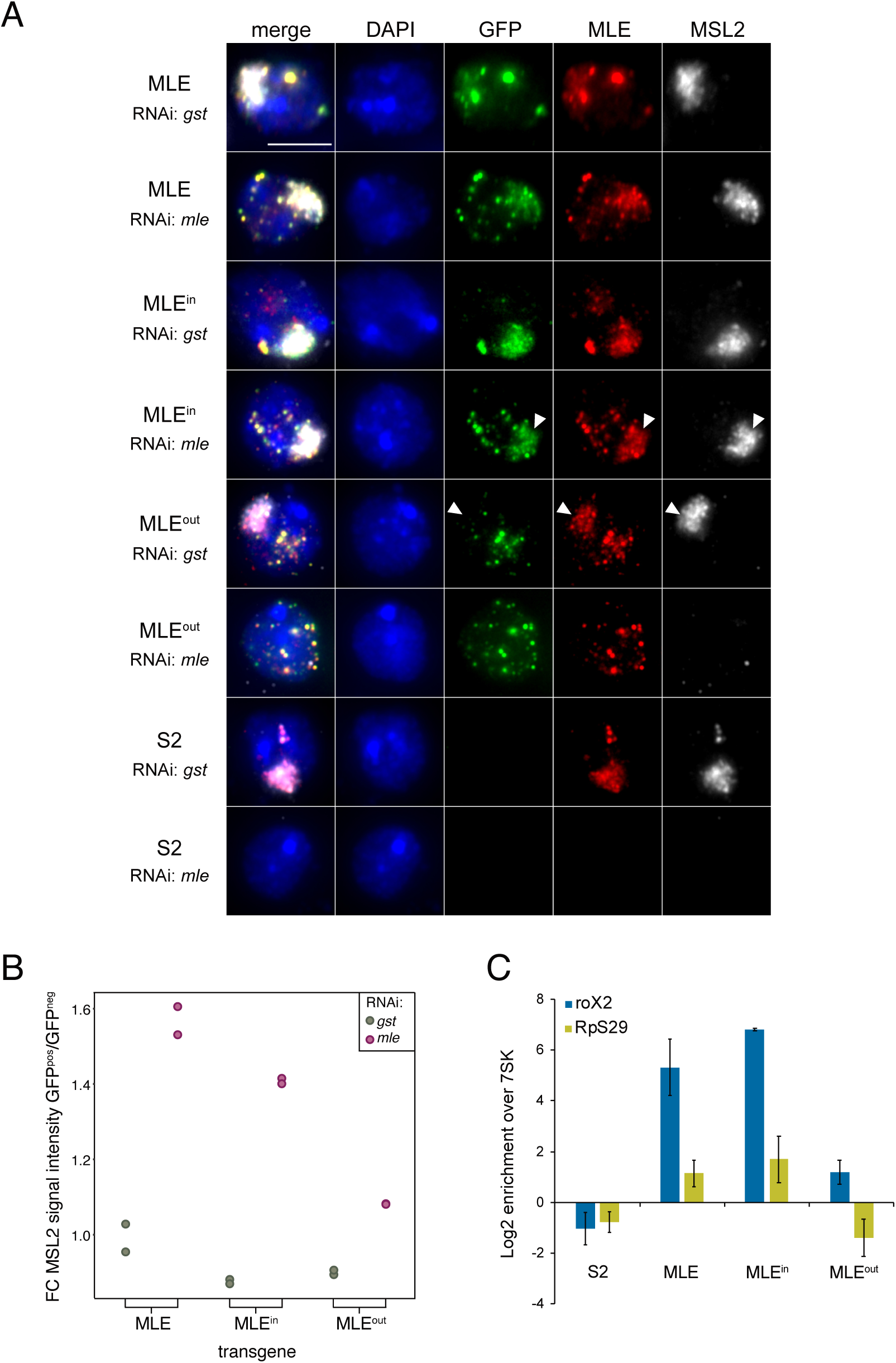
Restricting dsRBD2 conformation affects X chromosome localisation, territory formation and RNA binding *in vivo*. (A) Representative immunofluorescence images of S2 cells stably expressing RNAi-resistant MLE_fl_-eGFP. Cells were treated with control (*gst*) dsRNA (endogenous and transgenic MLE present) or with dsRNA targeting endogenous *mle* (endogenous MLE depleted and transgenic MLE present). Panel ‘S2’ shows immunofluorescence of non-transfected S2 control cells, which were treated the same way. DNA is shown in blue, transgenic MLE_fl_-eGFP in green, MLE in red and MSL2 in grey. Arrowheads mark X territories. Scale bar represents 5 µm. (B) Dot plot showing the immunofluorescence-based complementation assay from (A). Each dot represents the fold-change of the mean nuclear MSL2 signal between GFP-positive (expressed MLE_fl_-eGFP wild type or mutant transgene) and GFP-negative cells per biological replicate. The number of cells included in the quantification is given in Figure S7B. (C) RNA immunoprecipitation (RIP) of GFP-tagged MLE_fl_-wild type and mutants from stable S2 cells. ‘S2’ represents a GFP-RIP control using non-transfected S2 cell extract. Relative enrichment (IP/input) of roX2 and RpS29 transcripts was analyzed by RT-qPCR and is presented normalized to unbound 7SK. Error bars represent the standard deviation for four independent replicates.

MLE binds with high selectivity to tandem stem loop structures and polyA/U-rich ssRNA sequence motifs, which in combination are present in critical parts of roX2 (Ilik *et al*., 2017; Maenner *et al*., 2013; Müller *et al*., 2020). Thus, for recognition and unwinding of roX2 both dsRNA and ssRNA binding modes are required. We reasoned that the observed phenotype in MLE ^out^-eGFP expressing cells is the consequence of compromised roX2 recognition and incorporation into the MSL-DCC (Figure 6A-B, S7C). Therefore, we assessed physiological substrate binding in MLE_fl_-eGFP expressing cell lines by RNA immunoprecipitation (RIP) under native, non-crosslinked conditions. These experiments were performed in presence of endogenous MLE to maintain a stable pool of MLE substrates in the cells. GFP-tagged MLE variants were immunoprecipitated using an anti-GFP antibody and the fraction of bound RNA was determined by RT-qPCR using primers targeting either the major substrate roX2 or the recently identified substrate RpS29 (Ilik *et al*., 2017; Müller *et al*., 2020). RIP of the abundant non-coding 7SK or GAPDH mRNA were used for normalization and extracts from non-transfected S2 cells served as negative control. As expected, roX2 immunoprecipitated well with MLE ^WT^-eGFP and, even to a larger extent, with the MLE ^in^-eGFP mutant, which in cells is considered wild-type like (Figure 6C, S7D). The MLE ^out^-eGFP mutant, however, showed severely diminished roX2 binding, suggesting that disrupting dsRBD2-helicase module interactions causes helicase deficiency and limits faithful roX2 recognition and stable integration into the DCC (Figure 6C, S7D). This observation is further strengthened by the fact that the dsRBD2 mutants are immunoprecipitated at higher levels than MLE ^WT^-eGFP due to a larger fraction of mutant transgene-expressing cells (Figure S7E).

Because roX2 is rendered unstable in absence of functional MLE (Franke and Baker, 1999; Meller et al., 2000), we propose that the observed failure of MLE ^out^-eGFP to complement MLE loss is explained by a deficiency of roX2 incorporation into the DCC (requiring helicase activity), leading to degradation of the RNA. In addition to roX1 and roX2, MLE binds – presumably transiently – to a small subset of additional substrates that harbor similar secondary structure and sequence determinants (Ilik *et al*., 2017; Ilik *et al*., 2013; Müller *et al*., 2020). To test if other substrates are recognized in the same way, we quantified the enrichment of RpS29 with the MLE_fl_-eGFP transgenes. MLE ^WT^-eGFP and MLE ^in^-eGFP indeed enriched RpS29 from S2 cell extracts, albeit at lower levels than roX2, confirming our previous observation (Figure 6C, S7D) (Müller *et al*., 2020). In contrast, the MLE ^out^-eGFP mutant did not enrich RpS29, suggesting that a comparable recognition mode applies to at least two substrates of MLE, roX2 and RpS29.

## Discussion

RNA helicases are molecular motors that unwind double-stranded RNA. This simple statement contrasts the observed complexity of domain organization and functional specification of many such enzymes. MLE, for example, exhibits an exquisite substrate selectivity: preferred substrates combine RNA secondary structure (pairs of stem-loops with alternative base pairing potential) with small linear motifs that are exposed only upon unwinding (the U-rich roX-box) (Müller 2020). For the unwinding reaction to be productive, MLE needs to control the helicase reaction, to hold on to the remodeled RNA and finally transfer the remodeled roX to the DCC.

We imagine that the observed substrate selectivity and context-dependence of MLE helicase is mediated by its several accessory domains, in interesting analogy to the structurally related nucleosome remodeling ATPases, where multiple accessory domains install conditional autoinhibition, substrate-mediated activation and proof-reading (Clapier et al., 2017). However, despite progress in the structural characterization of the MLE helicase motor or it’s accessory dsRBD domains in isolation (Jagtap *et al*., 2019; Lv *et al*., 2019; Prabu *et al*., 2015) an integrated view of their cooperation had been lacking.

Our new high-resolution structures reveal dramatic conformational changes of the MLE helicase upon binding to substrate RNA, leading to conditional opening of the channel that threads the ssRNA products through the enzyme. Combined with previous studies these structures allow us now to propose a first model for the helicase cycle of MLE. Below, we highlight the salient features of the model, which is visualized in Figure 7.

**Figure 7:**
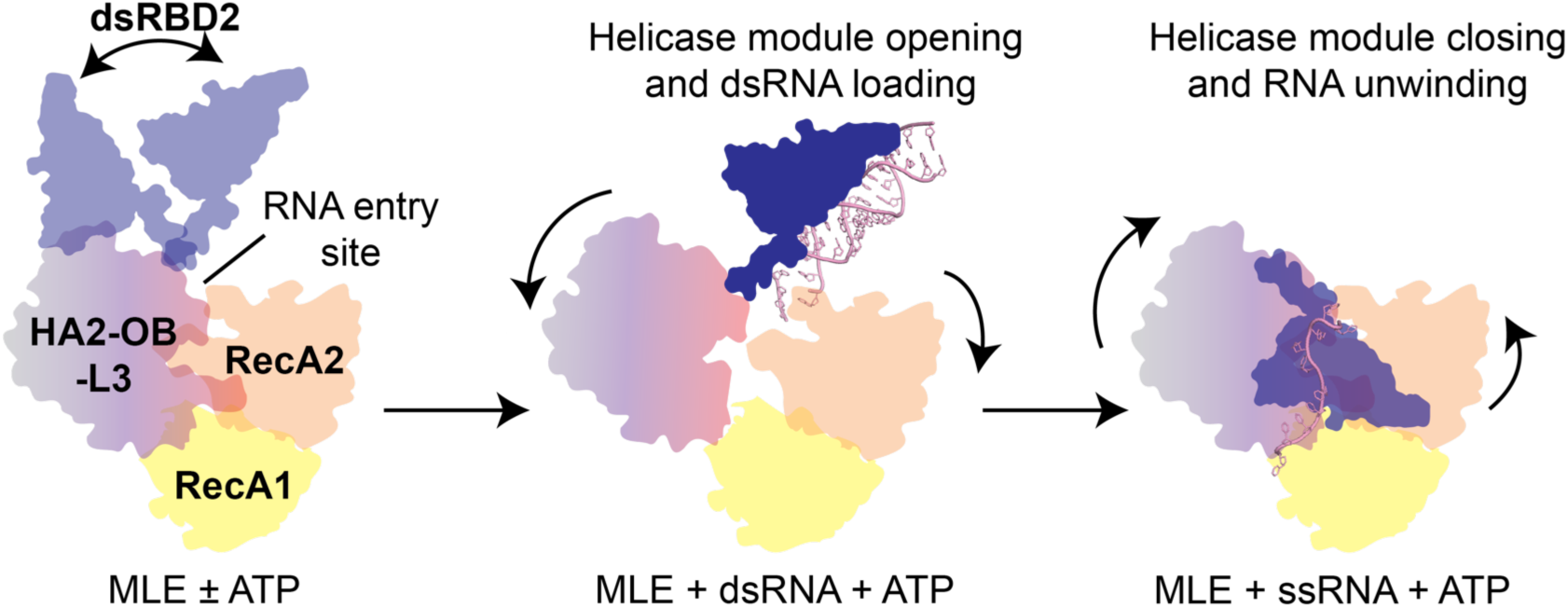
A model for cis regulation of RNA binding and helicase activity by accessory domains in MLE. dsRBD1 (not shown) and dsRBD2 domains remain flexible relative to each other and the helicase module in the apo and nucleotide bound states, allowing them to bind and retrieve dsRNAs. In the absence of RNA, the helicase module itself stays in a closed conformation, possibly to prevent binding of non-physiological RNA substrates. The limited linker length between dsRBD2 and L2 linker ensures productive binding of the helicase module to the 3’-end or to single stranded loops of dsRNA substrates. Recognition of the first few nucleotides from the single stranded RNA regions at the RNA entry site in the helicase module triggers the opening of the RNA binding tunnel: RecA2 and the HA2-OB-L3 module moves away from the tunnel thereby creating binding sites for the ssRNA within the tunnel. Since structural elements of HA2 and RecA2 domains occupy the positions in the middle of the RNA binding tunnel, the ssRNA is still not completely accommodated in the tunnel. For faithful ssRNA binding within the tunnel, dsRBD2 dissociates from the dsRNA and flips back onto the helicase module, thereby causing a movement of HA2 and RecA2 structural elements away from the ssRNA binding path. This would also force the helicase module into the RNA bound closed conformation allowing the helicase activity of MLE.

### Hallmarks of the model

In our model (Figure 7) the helicase module adopts a closed conformation, which prevents binding of the non-substrate RNA. Connected by flexible linkers, the two dsRBDs domains are free to sample the immediate vicinity for potential substrate RNAs. The defined length between dsRBD2 and L2 linker ensures productive binding of captured RNAs by the helicase module either at the 3’-end of dsRNA or in the single stranded loops. Recognition of first nucleotides from the unwound RNA at the RNA entry site in the helicase module induces movement of RecA2 domain and HA2-OB-L3 modules thus opening of the RNA binding tunnel. To completely free the tunnel, dsRBD2 must dissociate from the dsRNA and flip back onto the helicase module, pulling back aspects of HA2 and RecA2 domains that otherwise occlude the central part of the tunnel. This conformational change is predicted to force the helicase module into the RNA-bound closed conformation allowing helicase activity and stable binding of the ssRNA product.

### Accessory domains regulate access to the RNA tunnel in DExH helicases

The ssRNA binding site in MLE and other eukaryotic DExH/DEAH RNA helicases consists of a tunnel formed by RecA1, RecA2 and accessory domains. Although several structures of these helicases especially in the context of spliceosomes have been determined recently, they lack their dsRNA substrates. A structure of DExH helicase Hel308 from archaea has been determined in complex with dsRNA, however the domain architecture and arrangement between Hel308 and eukaryotic DExH/DEAH RNA helicases is very different (Buttner et al., 2007). The specific architecture of accessory domains relative to the helicase module undoubtedly contributes to substrate selectivity and regulation among helicases.

Previous DExH/DEAH RNA helicase structures have shown that binding of the ATP transition state analogue opens the ssRNA binding tunnel by rearranging C-terminal domains (Tauchert *et al*., 2017). In interesting contrast, we did not observe any significant conformational changes upon binding of ADP:AlF_4_ in MLE, suggesting a different, ATP-independent mechanism for opening up of the ssRNA binding tunnel (Figure S2C). Such a mechanism of a nucleotide-independent RNA tunnel opening in the helicase module has so far only been observed in case of the DEAH/RHA helicase DHX36, which unwinds G-quadruplexes (Chen et al., 2018). Our structure of MLE_ΔG_ in complex with dsRNA provides a first snapshot of how dsRNA enters into the RNA tunnel in this class of helicases: In addition to their role in direct RNA binding, HA2 and the OB-like domains function as a gate that needs to be opened by large domain rearrangements to regulate the entry of RNA into the tunnel.

### Selective binding of RNA by MLE could occur early during RNA recognition

In the structures of MLE in complex with ssRNAs, determined in this study and previously (Prabu *et al*., 2015), the first two nucleotides of ssRNA bend towards and interact with the RecA2 and dsRBD2 domains. This is different in the dsRNA-bound structure, where the first nucleotide which enters the core interacts with K1033 from the OB-like β_3_-β_4_ loop in the dsRNA bound structure (Figure 2B). Given the orientation of the RNA 3’-end of dsRNA in this structure and from its superposition with ssRNA bound structures, the RNA path of the 3’-end of dsRNA in the RNA tunnel would likely continue from the position corresponding to the 3^rd^ nucleotide in ssRNA bound structures (Figure S4B). This 3^rd^ nucleotide in ssRNA bound structures is recognized in a base-specific manner by residues from the RecA2 and OB-like domains (Prabu *et al*., 2015). Therefore, these accessory domains may be involved in a specificity checkpoint, selecting against binding of non-substrate RNAs. Previous mutations of H1032 and K1033 from the OB-like domain strongly reduced the affinity of MLE to not only ssRNA but also to dsRNA (Prabu *et al*., 2015). Our MLE structure in complex with the dsRNA provides a structural explanation for this, as K1033 directly contacts the 3’-end of the backbone phosphate of dsRNA (Figure 2B). Such a mechanism, where auxiliary domains shape the RNA preferences in helicases, has been recently been reported in the bacterial DExH helicase HrpB (Hausmann et al., 2020).

### Role of dsRBD2 in MLE beyond dsRNA recognition

Previously, the requirement of dsRBD2 for RNA binding and helicase activity of MLE was exclusively interpreted in the context of its role in dsRNA binding (Izzo *et al*., 2008; Jagtap *et al*., 2019; Lv *et al*., 2019; Prabu *et al*., 2015). We were now able to design structure-guided mutations which uncouple the contributions of dsRBD2 for dsRNA and ssRNA binding and its role in modulating the helicase activity. Remarkably, we found that association of dsRBD2 with the helicase module triggers the closed conformation of MLE by acting as a brace between RecA2 and the HA2-OB-L3 module and thus stabilizes binding of ssRNA in the core. This mode of action is distinct from its binding to dsRNA, where dsRBD2 must dissociate from the helicase module to allow it to open (Figure 3).

In MLE, the proposed gating mechanism is implemented through the conformational changes of dsRBD2, which works as an autoregulatory principle *in cis*. Recently, a similar helicase activation mechanism, albeit in *trans*, was shown for Prp43. Here, a G-patch-containing protein acts as a brace between RecA2 and the winged helix domains thereby tethering two mobile parts of the helicase together. In addition, *trans* activation by related G-patch proteins have been proposed for Prp16 and DHX35 (Bohnsack et al., 2021; Studer et al., 2020). These proteins, like many RNA helicases, need *trans* factors to boost their inherently poor helicase activity (Ozgur *et al*., 2015; Silverman et al., 2003; Sloan and Bohnsack, 2018). The comparison between different DExH helicases beautifully illustrates how evolution finds different solutions to a structural problem.

DExH helicases require single-stranded 3’ overhangs to engage with substrates (Jankowsky, 2011; Ozgur *et al*., 2015; Pyle, 2008). Curiously, MLE was shown to be able to bind and unwind RNA duplexes lacking overhangs (Prabu *et al*., 2015). Our structures now suggest a molecular explanation. dsRBD2 and the linker that connects it with the helicase module provide a continuous positively charged surface, which holds the dsRNA prior to entry into the helicase module (Figure S3G). Interactions between the linker and the helicase module restrict the flexibility of dsRBD2 thereby holding the dsRNA in a conformation primed for entry into the active site. Its limited length constrains the interaction of dsRBD2 towards the base of large RNA stems (Figure 2B).

In striking contrast to other helicases, such as Prp43, MLE shows a profound selectivity for uridine-rich sequences, which engage in base-specific contacts in the ssRNA channel. This raises the question about how such tightly bound ssRNA would be released during the helicase cycle and handed over to the DCC. Our structures now suggest a plausible mechanism: ssRNA can be released either stochastically or in a regulated manner when dsRBD2 detaches from the MLE helicase module, thereby forcing its opening. It is tempting to speculate that such a mechanism could be directly connected to the ‘handover’ of unwound roX RNA to MSL2, which has been shown to also bind the U-rich roX box (Müller *et al*., 2020).

### Potential role of dsRBD1 in mediating protein-protein interactions in MLE

Previously, only dsRBD2 was shown to interact with the MLE helicase module (Prabu *et al*., 2015). However, our NMR titration combined with CL-MS analysis and cryo-EM structures showed that dsRBD1 interacts transiently and non-specifically with the helicase module (Figure 4). dsRBD1 binds only weakly to RNA, which is further confirmed by our MLE_ΔG_ structure in complex with the dsRNA, where dsRBD1 is not visible in the cryo-EM density (Figure 2). dsRBD1 was shown to be required for the proper localization of MLE with the MSL complex at the X chromosome, but mutation of dsRBD1 RNA binding residues had minor effects on the localization of MLE with the MSL complex (Izzo *et al*., 2008; Jagtap *et al*., 2019; Lv *et al*., 2019). Our data support earlier suggestions of a potential role for dsRBD1 in mediating protein-protein interactions during DCC assembly. Within DCC, these factors could be the MSL1/MSL2 module which is sufficient for the recruitment of the MLE-roX2 complex to the DCC (Müller *et al*., 2020).

### Implications for human DHX9 mechanism and inhibition

Understanding the mechanics of MLE helicase function also sheds light on the function of its human orthologue, DHX9/ RNA helicase A (RHA). DXH9 and MLE show a similar domain organization with 51% sequence identity within structured domains. Conceivably, DHX9 is thus regulated by similar dsRBD2-mediated substrate gating mechanisms. DHX9 is a multi-functional protein with roles in transcription regulation, mRNA translation and miRNA processing (Anderson et al., 1998; Hartman et al., 2006; Nakajima et al., 1997; Robb and Rana, 2007). It is also involved in the replication of various viruses such as hepatitis C and HIV and directly interacts with inverted *Alu* transposon elements to suppress the RNA processing defects arising due to *Alu* transposon insertions (Aktas et al., 2017; Isken et al., 2003; Tang et al., 1997).

Considering its role in tumour cell maintenance, targeting of DHX9 could provide a novel chemotherapeutic approach (Gulliver et al., 2020; Lee et al., 2016). The previously identified inhibitor aurintricarboxylic acid is a general, non-specific inhibitor of protein-RNA interactions (Cencic et al., 2015; Gonzalez et al., 1980). Our study suggests that interfering with the interactions between dsRBD2 and the helicase module would provide a viable alternative strategy for drug development.

## Methods

### Protein expression and purification

N-terminal His_6_-tagged MLE_ΔG_ (1-1158), His_6_-tagged MLE helicase module (257-1158) and C-terminal flag-tagged MLE-full length and the mutant constructs and were cloned, expressed, and purified as described previously (Jagtap *et al*., 2019) in either pFastBac or pHsp70-eGFP plasmids. The MLE_ΔG_^in^ (with E195C and S633C mutations) and MLE_ΔG_^out^ (consisting of R590A, E602A, D603A, E605A, E608A D634A H746A, N747A, T754A, E790A, E835A, K1027A and R1057A mutations) mutants were generated by overlapping PCR and site-directed mutagenesis with primers containing the respective mutations in pFastBac MLE_ΔG_, pFastBac MLE_fl_ and pHsp70-MLE_fl_-eGFP plasmids. Around 700 bp of the original *mle* cDNA sequence in pHsp70-MLE_fl_-eGFP (wild-type and mutants, respectively) was substituted with a mutagenized, synthesized DNA sequence to generate RNAi-resistant *mle* constructs.

The proteins were expressed in SF21 insect cells using recombinant baculovirus. The cells were harvested after three days of infection and were resuspended in lysis buffer (20 mM HEPES, 500 mM NaCl, 20 mM imidazole, 2 mM β-mercaptoethanol, pH 7.5 with protease inhibitor and 10 units of Benzonase), lysed by sonication and spun down. The supernatant was loaded onto a HisTrap HP column (GE Healthcare) and eluted with an imidazole gradient in buffer containing 20 mM HEPES, 500 mM NaCl, 500 mM imidazole and 2 mM β-mercaptoethanol, pH 7.5. His_6_-tag was removed by addition of TEV protease and simultaneous dialysis into a low salt buffer (20 mM HEPES, 100 mM NaCl, 2 mM β-mercaptoethanol, pH 7.5) overnight at 4°C. The next day, the samples were applied to a second affinity chromatography using a HisTrap HP column, and the flow-through fraction was further loaded onto a HiTrap Heparin column (GE Healthcare) to remove bound bacterial RNA. The protein was eluted from the HiTrap Heparin column with a salt gradient in the elution buffer (20 mM HEPES, 1 M NaCl, 2 mM β-mercaptoethanol, pH 7.5). Fractions containing MLE were concentrated with Amicon Ultra centrifugal filters with a molecular weight cut-off of 30 kDa, and the sample was applied to size-exclusion chromatography on the HiLoad 16/60 Superdex S200 pg column (GE) in 20 mM HEPES, 150 mM NaCl, 5% Glycerol, 1 mM DTT, pH 7.5.

For NMR experiments, dsRBD1,2 domains were expressed and purified as described previously (Jagtap *et al*., 2019). Prior to NMR experiments, the MLE helicase module was dialysed in the NMR buffer containing 20 mM potassium phosphate, 200 mM NaCl, pH 6.5. To the final NMR samples, 10% D_2_O was added for the deuterium lock.

### In vitro transcription

Stem-loop 678 roX2 RNA (SL678) (5’-ACAAUAUGCAAUACAAUACAAUACAAGACAAAAAAAUGUGUCUUGGAACGCAACAUUGUACAAGUCGCAAUGCAAACUGAAGUCUUAAAAGACGUGUAAAAUGUUGCAAAUUAAGCAAAUAUAUAUGCAUAUAUGGGUAACGUUUUACGCGCCUUAACCAGUCAAAAUACAAAAUAAAUUGGUAAAUUUCAUAUAACUAGUGAAAUGUUAUACGAAACUUAACAAUUGCCAAAUAA-3’) was prepared by *in vitro* transcription using in-house produced T7 polymerase, rNTPs, and the RNA sequence cloned into pUC19 plasmid as template. The pUC19 DNA was linearised using EcoRI. After transcription, the reaction was cleaned by phenol/chloroform extraction. The RNA was purified by denaturing 6% urea PAGE and extracted from the gel by electro-elution. The final sample was concentrated and precipitated using ethanol. RNA pellet from the precipitate was dissolved in water and the RNA samples were stored frozen at -20°C.

### Electromobility shift assays (EMSAs)

EMSA experiments were performed with SL7 RNA (GUGUAAAAUGUUGCUAGCAAAUAUAUAUGCUAGUAACGUUUUACGCCCUCUUUCUUUCUU) purchased from Integrated DNA Technologies (IDT) or with in vitro transcribed SL678 RNA. To label the RNA with Cy-5, 100 pmols of RNA were incubated with 150 pmoles of pCp-Cy-5 (Jena Biosciences), 20 units of T4-RNA ligase (NEB), 1x T4-RNA ligase buffer, and 1 mM ATP overnight at 4°C. After labelling, RNA was purified by phenol-chloroform extraction and ethanol precipitation. Before EMSA experiments, the labelled RNA was heated to 94 °C for 2 min in water and snap-cooled on ice for 10 mins. EMSA reactions were carried with indicated amount of the protein mixed with 5 nM of Cy-5 labelled RNA substrate in 20 mM HEPES, 50 mM NaCl, 1 mM DTT, 10% glycerol and 0.005% IGEPAL CA-630, pH 7.5. The reactions were incubated for 30 mins at room temperature and were loaded on the native 6% polyacrylamide gel. The gel was imaged in the Typhoon FLA 900 imager (excitation 651 nm and emission 670 nm). The experiments were performed in duplicates.

### Fluorescence polarisation assays

Fluorescence polarization assays were carried out in 20 mM HEPES, 100 mM NaCl, 0.005% IGEPAL CA-630, pH 7.5 and with Fluorescein labelled SL678 or UUC RNA. In vitro transcribed SL678 roX2 or commercially purchased UUC RNAs was labelled at the 3’-end with Fluorescein (Qiu et al., 2015). An increasing protein concentration was incubated with 2.5 nM of RNA for 30 mins in Corning 384 well plate in 40 μL volume. The fluorescence polarisation was measured in BioTek Synergy 4 plate reader with excitation and emission of 485 and 528 nm, respectively. Data were plotted in Graph Prism v9 and were fit to the Sigmoidal 4PL equation (where X is log (concentration)) (*Y* = *Bottom* + 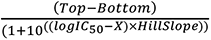 to calculate the IC_50_ values. All the experiments were performed in triplicates.

### Sample preparation for cryo-EM

Samples for cryo-EM grid preparation were prepared in the EM buffer (20 mM Tris pH 7.5, 50 mM NaCl, 1 mM DTT, transition state analogue (1 mM AlCl_3_, 10 mM NaF), 1 mM ADP, 2 mM MgCl_2_, 0.005 % triton-X 100, and 0.5% glycerol). RNAs used for the cryo-EM sample preparation were purchased from either Integrated DNA technologies (SL7 RNA, GUGUAAAAUGUUGCUAGCAAAUAUAUAUGC UAGUAACGUUUUACGCCCUCUUUCUUUCUU) or Biomers (UUC RNA, CCUCUUUCUUUC and U10-mer RNA). For preparation of the MLE-SL7 RNA complex, RNA was heated to 95°C for 2 mins and snap-cooled on ice. The MLE-RNA complex was prepared by mixing 4.7 μM MLE_ΔG_ with 5.2 μM (1.1 molar excess) RNA in the EM buffer. Before applying to the grid, the samples were incubated on ice for 30 mins. EM sample for MLE_ΔG_^apo^ was prepared without ADP and transition state analogue.

### Cryo-EM grid preparation and imaging

Quantifoil grids were plasma cleaned with 90% Argon, and 10% Oxygen plasma for 30 sec. For MLE_ΔG_^apo^ and MLE_ΔG_ samples, Quantifoil UltrAufoil grids (R 2/2, 200 mesh) and for MLE_ΔG_ in complex with RNA, Quantifoil holey carbon grids (R2/1, 200 mesh) were used. 3.5 μl of the sample was applied to the grids, and the grids were plunged into liquid ethane using Vitrobot Mark IV at 6°C and 100% humidity. Cryo-EM data were collected on an FEI Titan Krios microscope operated at 300 kV, equipped with a K3 detector operating in counting mode and a post-column Gatan Bioquantum energy filter. For MLE_ΔG_^apo^, MLE_ΔG_, MLE_ΔG_+U10 and MLE_ΔG_+UUC, 5313, 5894, 19545, 8880 movies were collected. In the case of MLE_ΔG_+SL7, datasets were collected in two microscopy sessions, with 8761 and 6742 movies recorded in each session. All datasets were recorded at a nominal magnification of 105,000x, corresponding to 0.822 Å/pixel at the specimen level. In the case of MLE_ΔG_+SL7 RNA samples, initial screening datasets indicated severe preferred orientation with only one dominant view as apparent from 2D and 3D processing. As we could not resolve this issue by adding additives in buffers and grid types with different surface treatments, cryo-EM datasets for the MLE_ΔG_+SL7 samples were collected with 30 degrees stage pretilt.

### Cryo-EM data processing

All movies were aligned with Patch motion correction, and CTF parameters were determined using Patch CTF in Cryosparc (Punjani et al., 2017). Particles were picked using Warp (Tegunov and Cramer, 2019). Particles were extracted from the CTF corrected micrographs using coordinates imported from Warp. Extracted particles were subjected to several rounds of 2D classification, and junk and low-resolution 2D classes were removed from further analysis. Selected particles from 2D classification were used to create multiple ab initio maps. The maps that showed high-resolution features were subjected to multiple rounds of heterogeneous refinement followed by homogeneous and non-uniform refinement.

### Model building

All cryo-EM structures were modelled by manual rigid-body fitting of the atomic models of individual domain of dsRBD2, RecA1 and RecA2 domains and the HA2-OB-L3 module derived from the crystal structure of MLE_core_ in complex with ssRNA in UCSF Chimera in the initial stage of model building (Pettersen et al., 2004; Prabu *et al*., 2015). The dsRNA in the MLE_ΔG_+SL7 structure was built by placing an ideal A-form helix of 19 bp into the EM-density of the RNA. The modelling was further completed by multiple iterative rounds of manual model building in Coot interrupted by real-space refinement in Phenix software suite (Adams et al., 2010; Emsley and Cowtan, 2004). The structures were analysed in UCSF Chimera, UCSF ChimeraX and PyMOL and the figure were made using UCSF ChimeraX and PyMOL (Goddard et al., 2018; Pettersen *et al*., 2004).

### NMR experiments

^1^H,^15^N HSQC NMR experiments for dsRBD1,2 were performed with 0.01 mM ^15^N labelled dsRBD1,2 on an Avance III Bruker NMR spectrometer equipped with a cryogenic probe head at 298 K and a magnetic field strength corresponding to a Larmor frequency of 800 MHz. ^15^N labelled dsRBD1,2 was added to an equimolar ratio of unlabelled MLE helicase module in the presence or absence of two-fold molar excess of U10 RNA and ATP transition state analogue (1 mM AlCl_3_, 10 mM NaF, 1 mM ADP and 2 mM MgCl_2_). Data were processed with nmrPipe/nmrDraw and were analysed in the CCPN analysis software with chemical shifts obtained from BMRB (ID: 34326) (Delaglio et al., 1995; Vranken et al., 2005).

NMR ATPase assays were carried out in 20 mM deuterated Tris, 100 mM NaCl, 5 mM MgCl_2_ and ± 2 mM TCEP and 1 mM ATP, pH 7.4 with 500 nM protein and 1 μM roX2 SL678 RNA for experiments carried out in the presence of RNA. 1D NMR experiments were performed on an Avance III Bruker NMR spectrometer equipped with a cryogenic probe head and a magnetic field strength corresponding to a Larmor frequency of 600 MHz and at 303 K. For each experiment, 32768 points in the direct dimension were recorded with eight scans, and for each sample, experiments were recorded as pseudo-2D experiments with data recorded every 60 seconds for a total of 120 mins. The data were processed in NMRPipe/NMRDraw, and the resulting 1D spectra were converted to text files using pipe2txt.tcl script from NMRPipe (Delaglio *et al*., 1995). The peaks corresponding to the H8 proton of ATP and ADP were integrated using custom scripts, and the individual peak intensities were divided by the total intensity of ATP+ADP peaks to obtain the percentage of ATP remaining or the percentage of ADP produced. The error bars were derived from the signal to noise ratio of the peaks. The data were then fit using one phase decay or one phase association equations to obtain the rate of ADP produced or ATP hydrolysed in GraphPad Prism 9.

### Cross-linking Mass spectrometry (CLMS)

Purified MLE_fl_-FLAG protein was centrifuged for 15 min at 21000 g at 4°C. MLE and RNA constructs of interest were incubated in a 1:2 molar ratio (0.7 µM MLE, 1.4 µM RNA) in MLE crosslinking buffer MXB-50 (20 mM HEPES pH 7.5, 50 mM NaCl, 2 mM MgCl_2_, 1 mM DTT) in presence of 1 mM Adenylyl-imidodiphosphate (AMP-PNP) and RNase inhibitor RNAsin (0.5 U, Promega) for 25 min at 4°C with head-over-end rotation. Samples were incubated with 1 mM Bis(sulfosuccinimidyl)suberate BS3 for 30 min at 30°C and 950 rpm. The crosslinking reaction was quenched by addition of 50 mM Tris-HCl/pH 8.0 and incubation for 15 min at 30°C and 950 rpm. A sample for Western blot analysis was taken to analyze the crosslinking degree of MLE.

Before tryptic digestion, 1 M urea was added to the samples to allow partial unfolding of the protein. Trypsin/Lys-C Mix, Mass Spec Grade was added in a 1:50 ratio to MLE. Tryptic digestion was carried out overnight at 37°C with 500 rpm agitation in presence of 1 mM DTT. Then 4.3 mM iodoacetamide was added and incubated for 30 min at 25°C, 500 rpm in the dark. Iodoacetamide was quenched by addition of 20 mM DTT and incubation for 10 min at 25°C and 500 rpm. Samples were acidified by the addition of 0.05% trifluoracetic acid (TFA), the pH was adjusted to 1. SDB-RPS stage tip material was equilibrated by washes with 100% acetonitrile (ACN), activation buffer (30% methanol, 0.2% TFA) and equilibration buffer (0.2% TFA). Trypsinized samples were loaded to the equilibrated stage tips and centrifuged for 15 min at 500 g. After subsequent washes with wash buffer (1% TFA in isopropanol) and equilibration buffer, peptides were eluted into low protein binding Eppendorf reaction tubes with freshly prepared elution buffer (80% acetonitrile, 1.25% ammonia). After vacuum drying of the samples at 45°C, peptides were resuspended in MS sample buffer (0.3% TFA, 2% ACN in MS grade H_2_O).

Mass spectrometry was essentially performed as described (Harrer et al., 2018). Briefly, for LC-MS/MS analysis samples were injected in an RSLCnano Ultimate 3000 system and either separated in a 15-cm analytical column (75 μm ID home-packed with ReproSil-Pur C18-AQ 2.4 μm) with a 50-min gradient from 5 to 60% acetonitrile in 0.1% formic acid or in a 25-cm analytical column (75 µm ID, 1.6 µm C18) with a 50-min gradient from 2 to 37% acetonitrile in 0.1% formic acid. The effluent from the HPLC was directly electrosprayed into a QexactiveHF (Thermo) operated in data dependent mode to automatically switch between full scan MS and MS/MS acquisition. Survey full scan MS spectra (from m/z 375–1600) were acquired with resolution R=60,000 at m/z 400 (AGC target of 3x10^6^). The ten most intense peptide ions with charge states between 3 and 5 were sequentially isolated to a target value of 1x10^5^, and fragmented at 27% normalized collision energy. Typical mass spectrometric conditions were: spray voltage, 1.5 kV; no sheath and auxiliary gas flow; heated capillary temperature, 250°C; ion selection threshold, 33.000 counts.

### CLMS data analysis

The raw data files were first converted by the proteome discoverer 2.2 (Thermo scientific) xlinkx workflow for crosslink detection into the .mgf file format. Next, the .mgf files were analyzed by crossfinder (Forne et al., 2012; Mueller-Planitz, 2015) applying the following filter parameters for identification of cross-linking candidates: False-discovery rate (FDR) ≤0.05, number of fragment ions per spectrum ≥4, number of fragment ions per peptide ≥2, fractional intensity of assigned MS2 peaks ≥0.05, relative filter score: 95. Crosslinks were visualized using the xvis web browser for arch plots (Grimm et al., 2015) (https://xvis.genzentrum.lmu.de).

### Circular dichroism spectroscopy

The circular dichroism spectra of MLE_ΔG_ and its mutants were recorded between 260 nm and 190 nm in a 0.2 mm cuvette, at 20°C at a concentration of 1 mg/ml in 20 mM Tris pH 7.5, 150 mM NaCl, 2% glycerol and ± 2 mM TCEP. The spectra were recorded using a Jasco J-815 CD spectrometer with a 50 nm/min scan speed, digital integration time of 1 sec with a 1 nm bandwidth and 10 accumulations. The data were plotted using GraphPad Prism 9.

### Size exclusion chromatography – multi-angle light scattering (SEC-MALS)

50 μl of MLE_ΔG_ or the mutants were injected onto a Superdex 200 5/150 GL gel-filtration column (Cytiva) in 20 mM Tris pH 7.5, 100 mM NaCl ± 2 mM TCEP, at a flow rate of 0.3 ml/min and at room temperature. The column was connected to a MiniDAWN MALS detector and Optilab differential refractive index detector (Wyatt Technology). Data were analyzed using the Astra 7 software (Wyatt Technology) and was plotted using GraphPad Prism 9.

### Protein thermal stability measurements

The thermal stability of the proteins was determined using Prometheus NT.48 nanoDSF instrument from NanoTemper Technologies. The protein was incubated at 1 mg/ml in 20 mM Tris, 100 mM NaCl ± 2 mM TCEP, pH 7.5 for 30 mins at room temperature and was loaded into the standard capillaries. The assay measured the tryptophan fluorescence at 330 nm and 350 nm from 20 - 90°C with a 1°C/min increase in temperature. The data were acquired and analysed with the PR. ThermControl v2.1.2 software provided with the instrument.

### Real-time fluorescence RNA helicase assay

Real-time fluorescence RNA helicase assays were carried out using a slight modification of the protocol described previously (Tani et al., 2010). Briefly, the assays were carried out in 20 mM Tris, 100 mM NaCl, 5 mM MgCl_2_, 0.005% IGEPAL CA-630, pH 7.4 and ± 2 mM TCEP and RNA probe with BHQ-1 and 6-FAM quencher-dye pair. Before the assembly of the reaction, the SL7-up RNA strand containing BHQ-1 at the 3’ end (GUGUAAAAUGUUGCUAGCA-BHQ1, Biomers) and SL7-down RNA strand containing 6-FAM at the 5’ end (6-FAM-UGCUAGUAACGUUUUACGCCCUCUUUCUUUCUU, Biomers) were mixed in equimolar ratio at 500 nM concentration in 20 mM Tris, 100 mM NaCl, pH 7.4, heated to 95°C for 1 min and then slowly cooled down to 4°C. Unlabelled SL7-up strand at a concentration of 250 nM was used as a competitor to prevent rehybridization of the BHQ-1-6-FAM RNA probe upon separation by the helicase. Reactions without protein were used as a control. The assays were carried out in triplicates in 40 μL reaction volume with 50 nM protein and 50 nM RNA probe in Corning 384 well plate. The fluorescent intensity of the fluorescein was measured in the BioTek Synergy 4 plate reader (excitation at 485 nm and emission at 528 nm wavelength) every 15 s at 30°C. 1 mM ATP was added to the reaction mix after initially monitoring the reaction for 3 mins. Reactions were monitored for 60 mins in total. Data were fit in GraphPad Prism 9 using plateau followed by two phase association equation 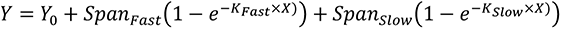, where K_Fast_ and K_slow_ are the rate constant for the fast and slow phase of the reaction given by *Span_Fast_* == (*Plateau* − *Y*_0_) × *Percent_Fast_* × 0.1 and *Span_Slow_* == (*Plateau* − *Y*_0_) × (100 − *Percent_Fast_*) respectively.

### Cell lines and culture conditions

*Drosophila melanogaster* male S2 cells (subclone L2-4, Patrick Heun, Edinburgh) were cultured in Schneider’s *Drosophila* medium (Gibco) supplemented with 10% fetal calf serum (Sigma-Aldrich) and penicillin-streptomycin at 26°C. Cell lines were tested negative for mycoplasma.

Stable S2 cell lines expressing wild type or mutant MLE_fl_ fused to C-terminal GFP were generated as described with minor modifications (Prabu *et al*., 2015). Briefly, 500 ng pHsp70-MLE_fl_-eGFP wild type or mutant plasmid was co-transfected with 25 ng plasmid encoding a blasticidin resistance gene using the Effectene transfection reagent (Qiagen). Stable MLE_fl_-eGFP expressing clones were selected in complete medium containing 25 ng/ml blasticidin for a duration of three weeks, followed by recovery in complete medium lacking blasticidin for another week. In the following, cell lines were cultured in complete medium lacking blasticidin.

### *In vivo* RNA immunoprecipitation (RIP)

Native RNA immunoprecipitation (RIP) of MLE_fl_-eGFP and mutant derivatives was performed as described in with modifications (Prabu *et al*., 2015). For each replicate, 0.7 x 10^8^ exponentially growing S2 cells expressing wild-type or mutant MLE_fl_-eGFP were collected by centrifugation (220 x g, 5 min), washed once with PBS and flash frozen in liquid nitrogen. Non-transfected S2 cells served as control and were treated likewise. Cell pellets were thawed on ice and resuspended in 700 µl of cold lysis buffer (20 mM HEPES-NaOH pH 7.6, 125 mM NaCl, 0.05% SDS, 0.25% sodium deoxycholate, 0.5% NP40, 1.5 mM MgCl_2_, 0.25 mM DTT) supplemented with 0.05 U/µl RNase-free recombinant DNase I (Roche), 0.4 U/µl RNasin (Promega) and 1x Complete EDTA-free protease inhibitor (Roche). The lysate was incubated for 20 minutes on ice with 5 seconds vortexing every 5 min and cleared by centrifugation (21,000 x g, 30 min, 4°C). 1.5% of the supernatant was kept on ice as input material for RNA extraction and Western blot, respectively. Per RIP, 30 µl GFP-Trap agarose beads (proteintech) were blocked with 2% (w/v) BSA and 0.1 mg/ml yeast tRNA (Sigma-Aldrich) in lysis buffer for 1 h at 4°C to minimize non-specific interactions. The beads were washed once in lysis buffer, mixed with the remaining supernatant and incubated at 4°C for 2 hours on a rotating wheel. Beads were washed with RIP-100, RIP-250, and RIP-100 buffer for 3 minutes each at 4°C (25 mM HEPES-NaOH pH 7.6, 0.05% NP40, 3 mM MgCl_2_ with 100 mM NaCl and 250 mM NaCl, respectively). RNA was extracted of 75% of the bead material using Proteinase K (100 µg in lysis buffer with 0.5 % SDS; 55°C for 45 minutes), phenol-chloroform extraction and ethanol precipitation in presence of 20 µg glycogen (Roche). Input material (1.5%) was treated equally. RNA pellets were resuspended in 20 µl RNase-free water. RNA input and IP material was analyzed by RT-qPCR using the SuperScript III First-Strand Synthesis System (Thermo Fisher) and Fast SYBR Green master mix (Applied Biosystems) with primers specific for roX2, RpS29, 7SK and GAPDH (Key Resources Table). roX2 and RpS29 RNA enrichment of MLE_fl_-eGFP and its mutants was calculated as IP/Input and normalized to unbound 7SK or GAPDH RNA. Western blot analysis of 1.5% input and 25% bead material was performed with antibodies against GFP and Lamin.

### RNAi interference and immunocytochemistry

RNA interference of target genes *mle* and *gst* was essentially performed as described (Maiato et al., 2003). Double-stranded RNA fragments (dsRNA) were generated using the MEGAscript T7 transcription kit (Thermo Fisher) using PCR-amplified DNA templates (for PCR primers, see Key Resources Table). RNA was precipitated using lithium chloride according to the manufacturer’s instructions. RNA pellets were resuspended in RNase-free water and annealed to dsRNA by incubation at 85°C for 10 min followed by slowly cooling down to 20°C.

For RNAi treatment, 1.5 x 10^6^ MLE_fl_-eGFP (wild type and mutant) expressing S2 cells and non-transfected S2 cells, respectively, were seeded in 6-well plates and supplemented with 10 micrograms of GST or MLE dsRNA. Cells were incubated with dsRNA for 7 days at 26°C. RNAi efficiency was controlled by Western blot analysis of 1 x 10^6^ cells with primary antibodies against MLE and Lamin.

Immunostaining with mouse anti-GFP, rat anti-MLE and rabbit anti-MSL2 primary antibodies was performed according to (Thomae et al., 2013). RNAi-treated cells were settled and fixed with PBS/3.7% paraformaldehyde for 10 min at room temperature, permeabilized with PBS/0.25% Triton X-100 for 6 min on ice and blocked with 3% BSA/PBS for 1h at room temperature. Cells on coverslips were incubated overnight at 4°C with primary antibodies. Following two washes with PBS/0.1% Triton X-100, fluorophore-coupled secondary antibodies donkey anti-mouse-Alexa488, donkey anti-rat-Cy3 and donkey anti-rabbit-Alexa647 were added for 1 h at room temperature. DNA was counterstained with DAPI. After PBS/0.1% Triton X-100 and PBS washes, cells were mounted in VECTASHIELD (Vector Laboratories).

### Fluorescence Microscopy

Fluorescence images were recorded at the core facility bioimaging of the Biomedical Center with a Leica Thunder Imager 3D Live Cell TIRF based on a DMi8 stand, equipped with a Leica DFC9000 GT sCMOS camera with 2048x2048 pixels and a Leica LED5 fluorescence excitation source with individually switchable LEDs for specific excitation. DAPI, GFP/Alexa Fluor 488 (MLE-GFP), Cy3 (endogenous and recombinant MLE) and Alexa647 (MSL2) signals were recorded with a Quad-Band filter cube and an additional emission filter wheel in the emission beam path to avoid tunnel crosstalk. Image stacks of 7 planes with a step size of 1 µm were recorded with a HC PL APO 100x/1.47 oil CORR TIRF objective at a pixel size of 65 nm.

Image processing, montage assembly and quantification was done in Fiji (Schindelin et al., 2012). Images shown are maximum intensity projections of z-stacks. Images were resized by a factor of 4 without interpolation followed by gaussian filtering with a radius of 2 pixels. Representative cells are shown for each cell line and RNAi condition.

Mean nuclear fluorescence quantification was done on maximum intensity projections of raw images. The macro code is available on request. Briefly: nuclei were segmented on median filtered DAPI and MSL2 sum images using Otsu dark auto thresholding upon rolling ball background subtraction. Clumped objects were separated with a Watershed algorithm. Objects (nuclei) were included in a size range of 12 – 60 sq µm and a circularity of 0.6 – 1.00. Objects at image edges were excluded. The ratio of the MSL2 mean nuclear signal between MLE-GFP positive and negative cells of two independent biological replicates was calculated and plotted using R-Studio.

## Data Availability

The MLE_ΔG_^apo^, MLE_ΔG_, MLE_ΔG_+U10, MLE_ΔG_+UUC and MLE_ΔG_+SL7 structures were submitted to the PDB under accession codes 8B9L, 8B9J, 8B9G, 8B9I and 8B9K and the cryo-EM density maps were submitted to the EMDB under the accession codes 15935, 15933, 15931, 15932 and 15934 respectively.

## Acknowledgements

We acknowledge support from Felix Weis and Wim Hagen for cryo-EM data collection and Bernd Simon for NMR data collection. We thank Raffaella Villa for providing RNAi-resistant *mle* cDNA and Silke Krause for technical support. We acknowledge the Protein Analytics Unit at the Biomedical Center, LMU Munich, for providing equipment, services, expertise and assistance with data analysis. The work was supported by EU Marie Curie Actions Cofund EIPOD fellowship to PKAJ, Deutsche Forschungsgemeinschaft (DFG) through grant Be1140/9-1 to PBB and an Emmy-Noether Fellowship to JH (HE7291).

## Author Contributions

Conceptualization: P.K.A.J., J.H.; Funding acquisition: J.H., P.B.B.; Investigation: P.K.A.J., M.M., A.K., A.W.T., K.L.; Software: P.K.A.J., A.W.T.; Supervision: J.H., P.B.B, M.B.; Writing original draft: P.K.A.J., J.H., P.B.B., M.M.; Writing-review and editing: P.K.A.J., M.M., A.K., A.W.T., K.L., M.B., P.B.B., J.H.

---

**Figure S1:**
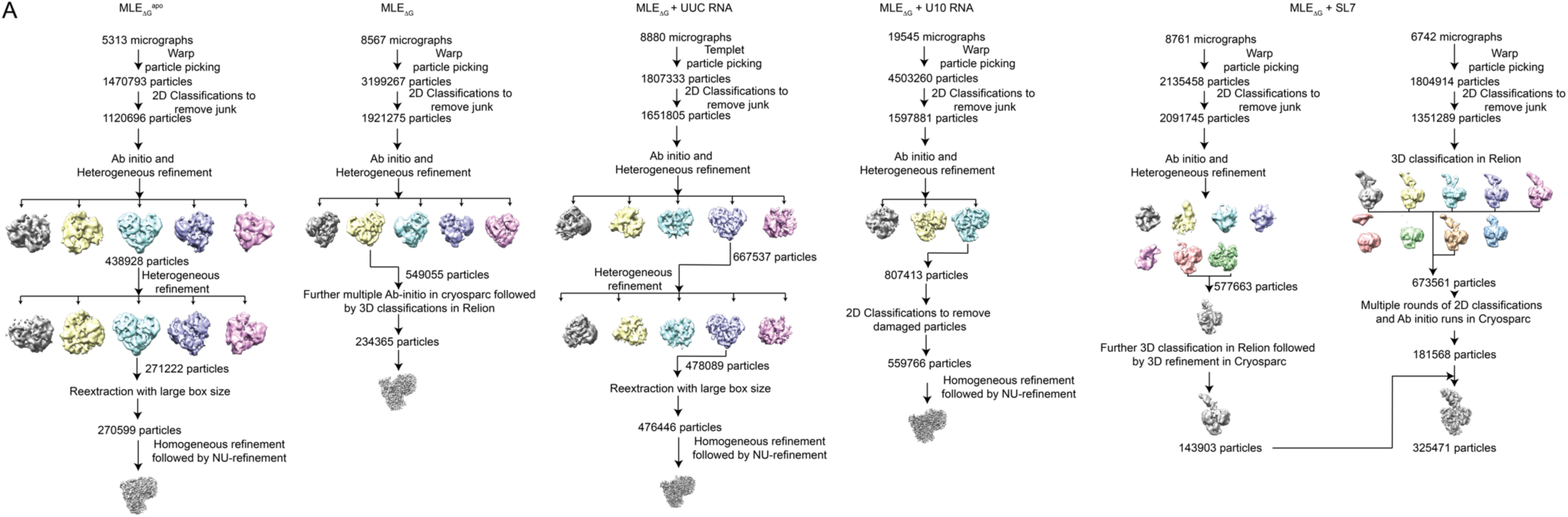
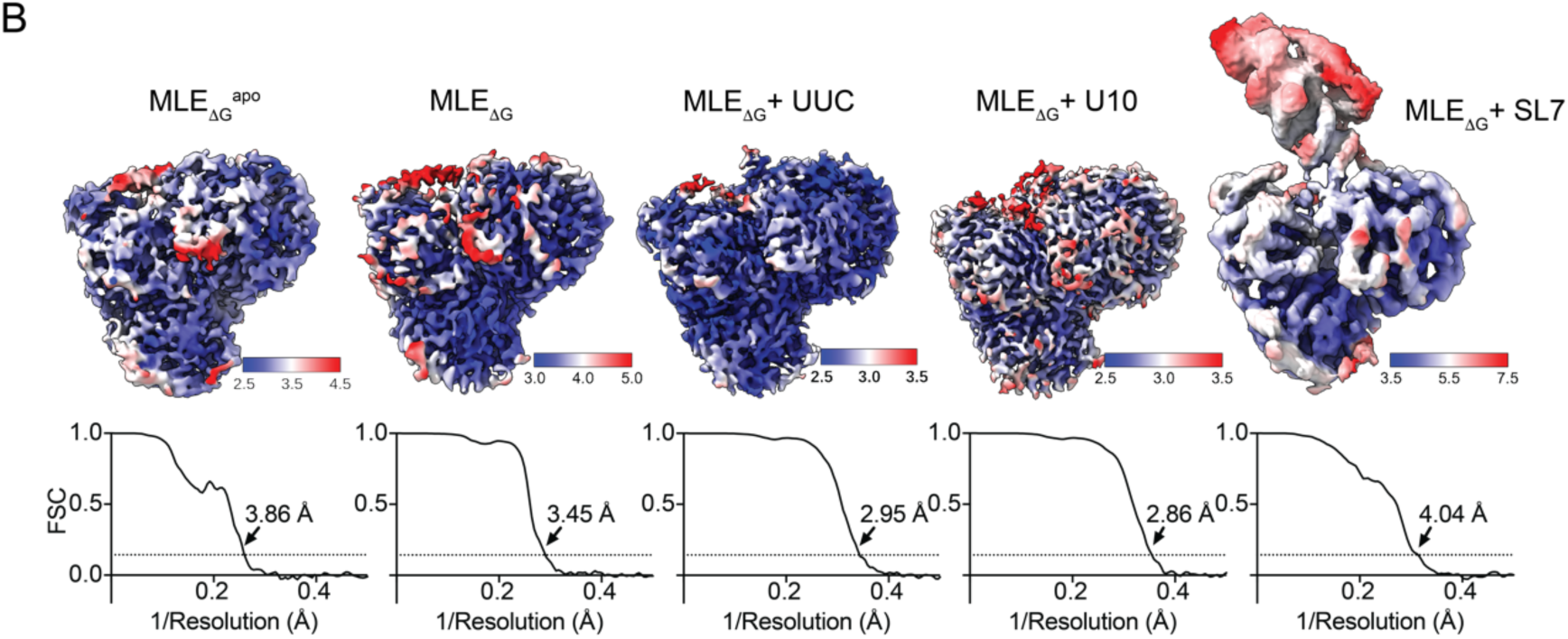
Cryo-EM processing and local resolution plotted on the EM density maps. (A) Cryo-EM processing pipeline used for all the structures determined for MLE_ΔG_ in the apo-state and in complex with ADP:AlF_4_, ss and dsRNA. (B) Local resolution and Fourier shell correlation curves along with resolutions (with 0.143 cut-off) for MLE structures.

**Figure S2:**
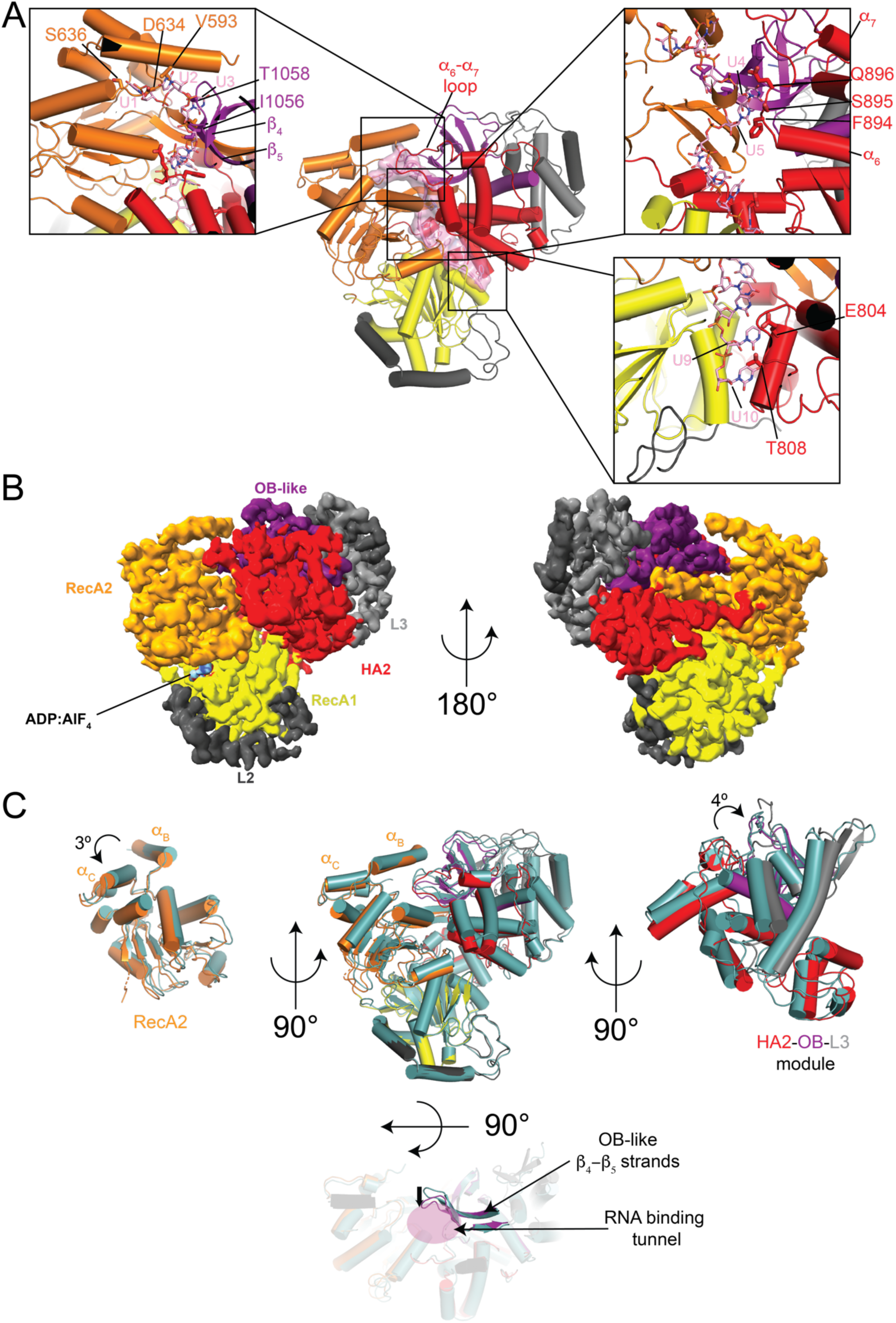
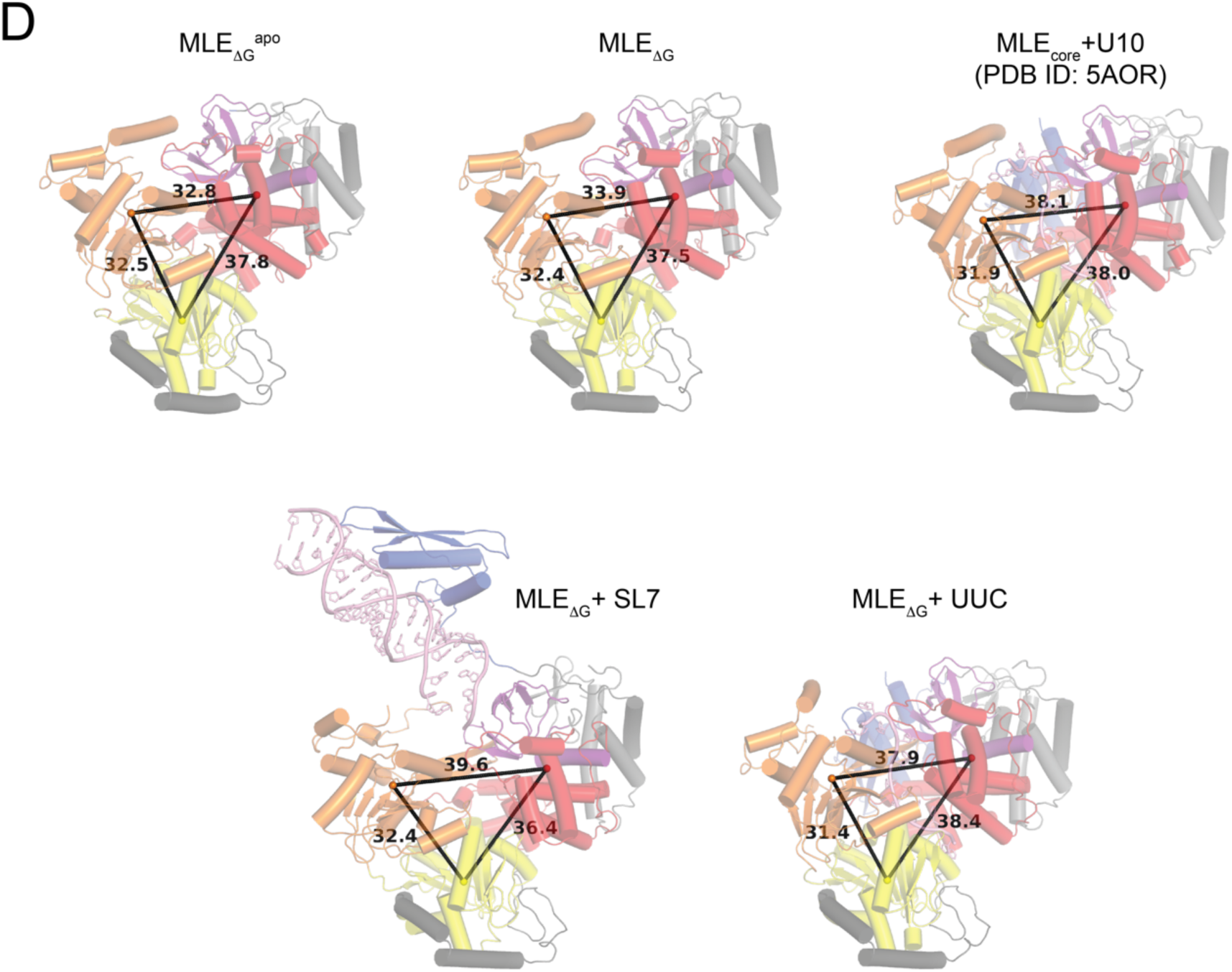
MLE helicase module exists in a closed conformation in the absence of RNA. (A) Superposition of U10 RNA from MLE_core_+U10 on to the MLE_ΔG_^apo^ structure. Zoom-in panels showing the clashes between the RNA nucleotides and the RecA2 and OB like domains at the entry, middle and exit tunnel of RNA. (B) Cryo-EM density map for MLE_ΔG_. (C) Structural superposition of MLE_ΔG_ (with respective domain colours) and MLE_ΔG_^apo^ (light teal) showing rotations in RecA2 domain and HA2-OB-L3 module. The inward movement of the β_4_-β_5_ sheet from OB-like domain, into the RNA binding tunnel, is shown with a black arrow in the top-view. (D) Centre of mass for RecA1 (yellow sphere), RecA2 (orange sphere) and HA2-OB-L3 module (red sphere) is shown along with the distances between them (in Å) for MLE structures.

**Figure S3:**
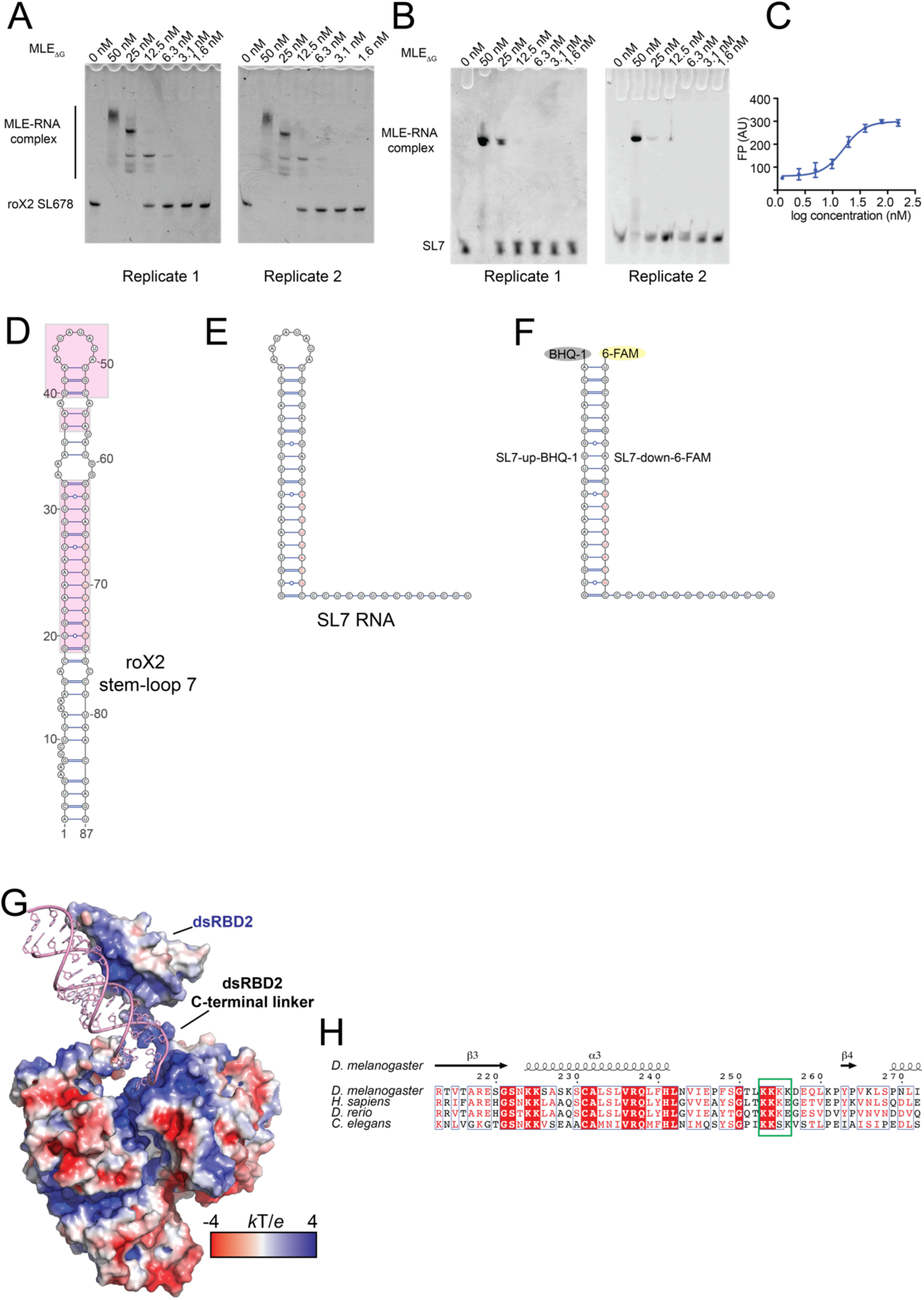
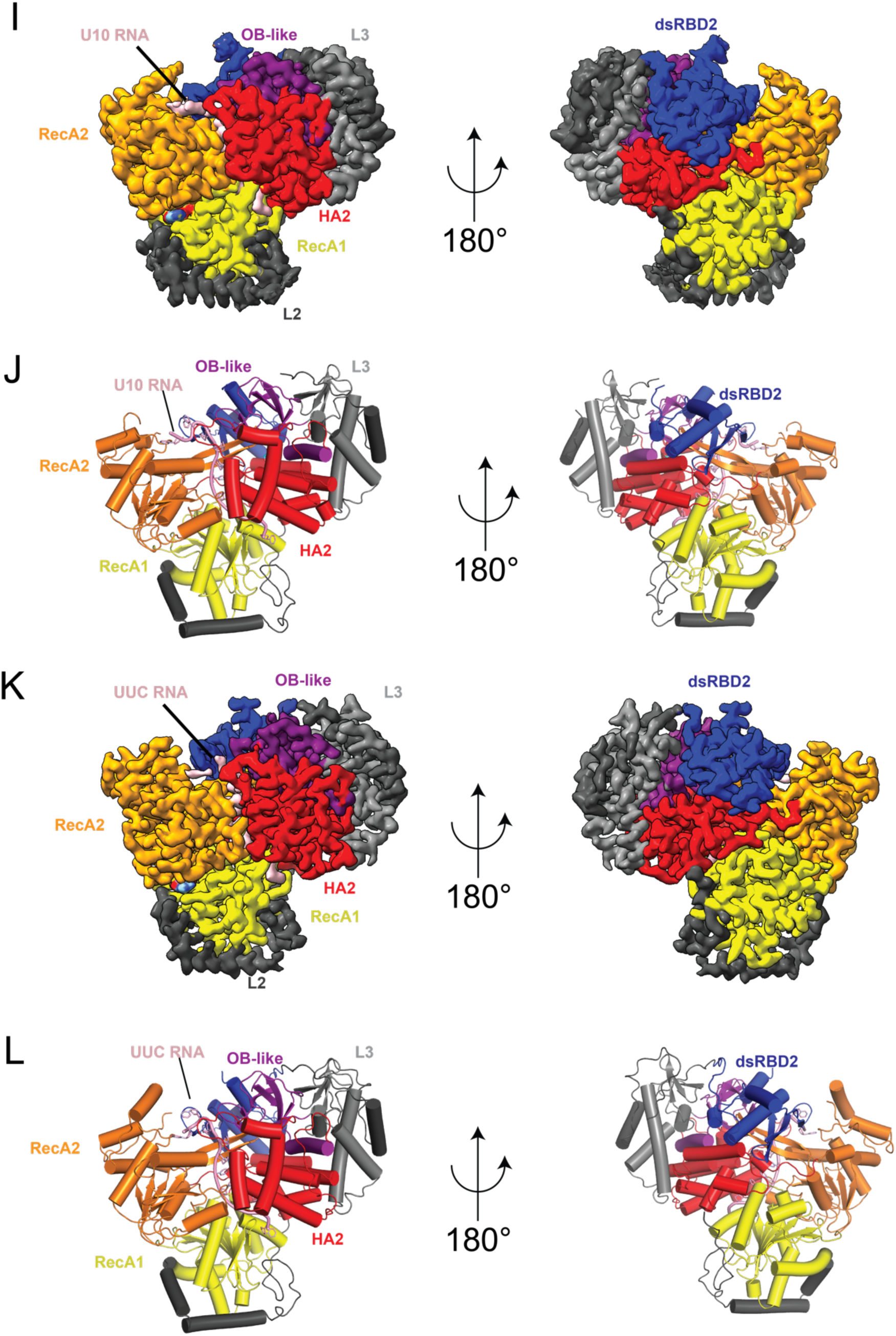
Structures of MLE_ΔG_ in complex with ds and ssRNA. EMSA gel showing binding of MLE_ΔG_ to (A) roX2 SL678 RNA and (B) SL7 RNA used in cryo-EM. (C) FP assay curve for the binding of SL7 dsRNA to MLE_ΔG_. (D) RNA secondary structure of roX2 SL7 RNA. The pink boxes highlight the regions used to design the SL7 RNA used for cryo-EM studies. (E) Secondary structure of the SL7 RNA and (F) RNA used for the real-time helicase assay. (G) Electrostatic potential plotted on the surface of the MLE_ΔG_+ SL7 structure. (H) Sequence alignment of the region between dsRBD2 and linker L2 reveals conservation of lysines 253-255. (I, J) Cryo-EM map and MLE_ΔG_ structure in complex with U10 ssRNA and ADP:AlF_4_ and (K, L) Cryo-EM map and MLE_ΔG_ structure in complex with UUC ssRNA and ADP:AlF_4_.

**Figure S4:**
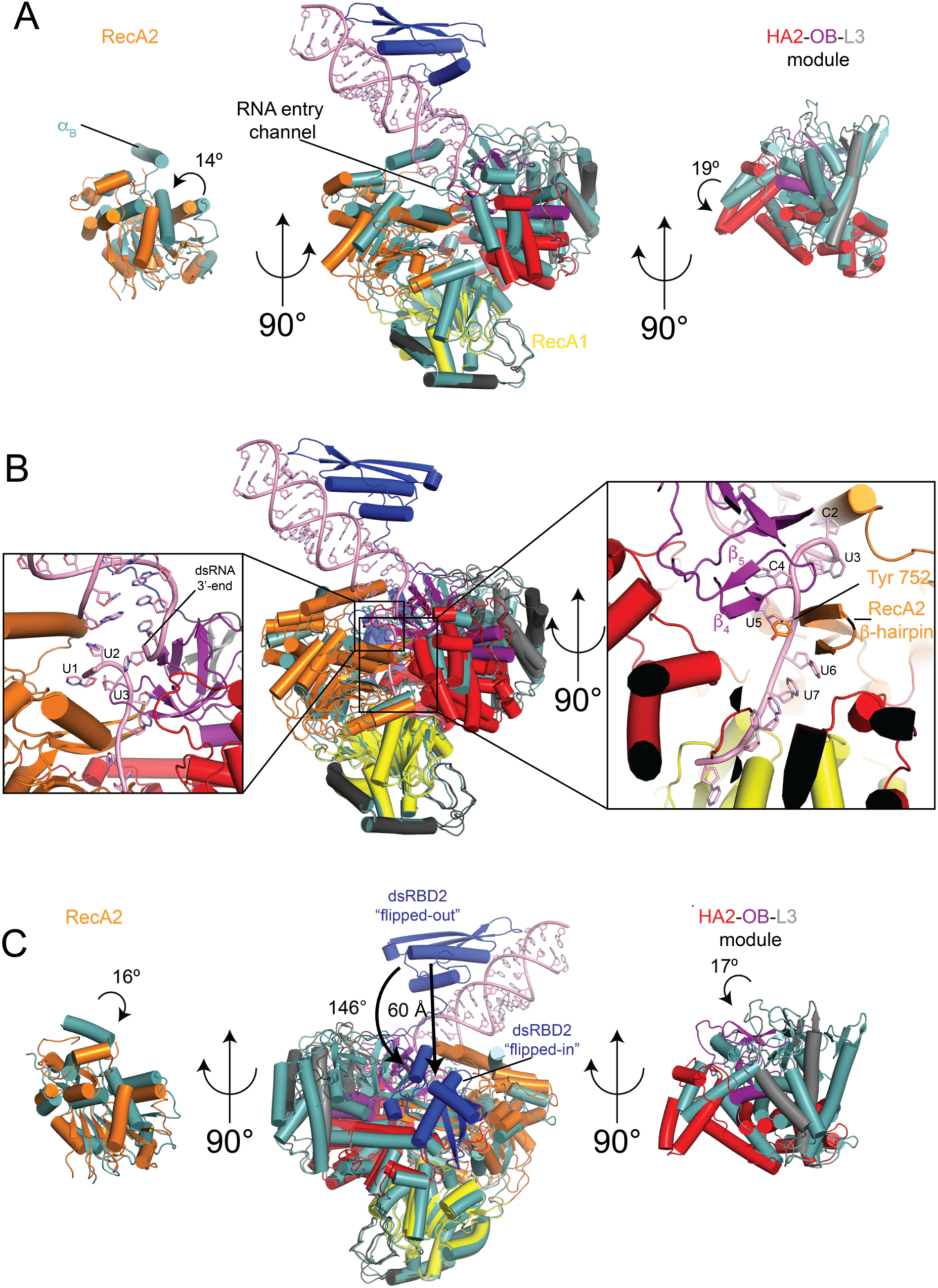
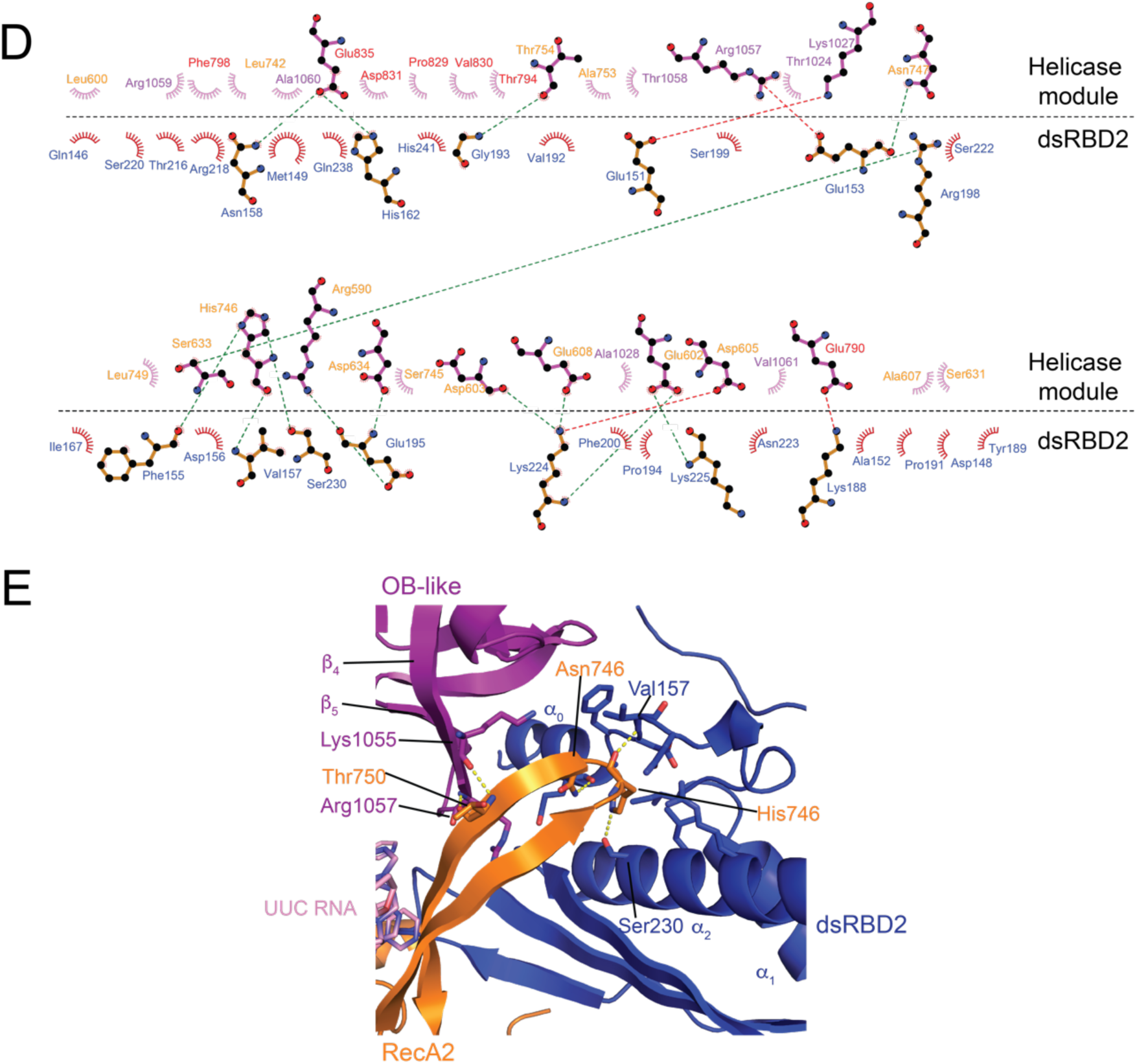
Structural changes and interdomain contacts in MLE upon binding to ds and ssRNA. (A) Structural superposition of MLE_ΔG_+SL7 (with respective domains colours) and MLE_ΔG_ (light teal) structures. For clarity, helices are shown as cylinders and ADP:AlF_4_ is not shown. Rotations of RecA2 and HA2-OB-L3 modules in the MLE_ΔG_+SL7 structure compared to the MLE_ΔG_ structure are shown in side view. (B) Structural superposition of MLE_ΔG_+UUC (with respective domains colours) and MLE_ΔG_+SL7 (light teal). Zoom-in inset shows the clashes between the UUC ssRNA and the dsRNA (SL7) bound MLE_ΔG_ structure which prevents the binding of UUC RNA extension from SL7 into the helicase module of MLE. (C) Structural superposition of MLE_ΔG_+SL7 (with respective domains colours) and MLE_ΔG_+UUC (dsRBD2 in blue, rest of the protein in light teal) structures. Structural changes in the dsRBD2, RecA2 domains and HA2-OB-L3 module are shown. (D) Interaction between dsRBD2 and helicase module plotted using Ligplot. Polar interactions are shown using green dashed lines. (E) Interactions of dsRBD2 in the MLE+UUC structure with the RecA1 β-hairpin are shown. α_0_ helix from dsRBD2 forms tight hydrophobic interactions with the OB-like domain thus pulling the OB-like β_4_-β_5_ strands out of the ssRNA path. In addition, the His746 side chain of RecA2 inserts itself into a shallow pocket formed by dsRBD2 α_0_-α_1_-α_2_ helices and forms hydrogen bonds with the backbone carbonyl of Val157 and the side chain of Ser230, along with several hydrophobic interactions. Asn747 from the RecA2 β-hairpin’s tip forms a hydrogen bond with the backbone carbonyl of Glu153 of dsRBD2. These interactions help to pull the RecA2 β-hairpin out of the ssRNA path within the RNA tunnel. Due to these conformational rearrangements of the HA2-OB-L3 module and the RecA2 domain, Thr750 from the RecA2 β-hairpin-1 forms several hydrogen bonds with Lys1055 and Arg1057 from the OB-like domain, thereby stabilising the OB-like β_4_-β_5_ strands at the RNA binding tunnel and further holding the RecA2 domain and the HA2-OB-L3 modules together.

**Figure S5:**
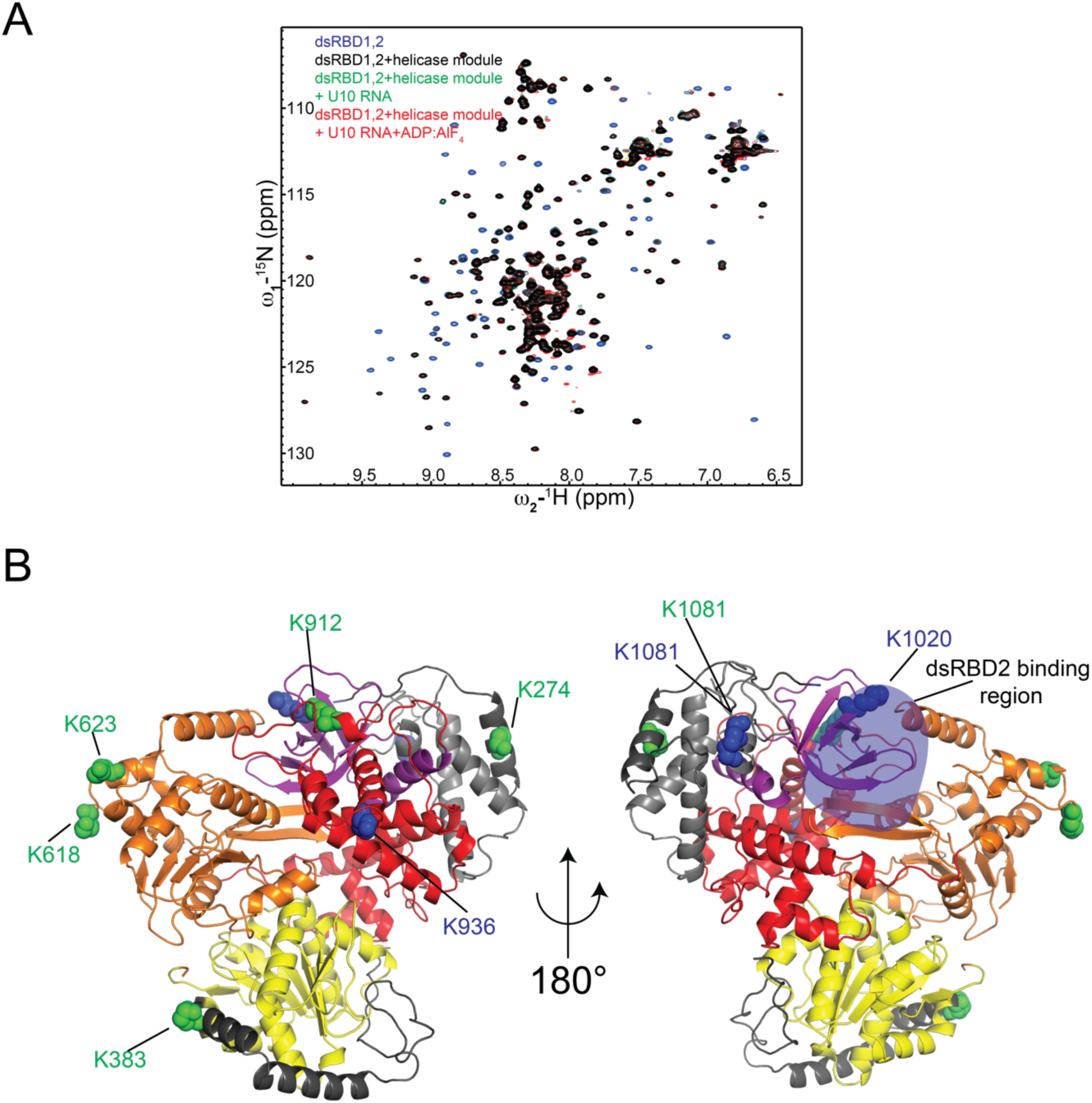
Structural characterization of interaction between dsRBD1,2 and helicase module of MLE. (A) Superposition of ^1^H,^15^N HSQC NMR titration spectrum of dsRBD1,2 free and bound to equimolar amount of MLE helicase module in the presence and absence of ss U10 RNA and ADP:AlF_4_. (B) Lysines in the helicase module which crosslink to dsRBD1 (shown in green) and dsRBD2 (shown in blue) are shown. K1081 crosslinks to both dsRBD1 and dsRBD2. For simplicity, lysines which crosslink to the linker between dsRBD1,2 are omitted.

**Figure S6:**
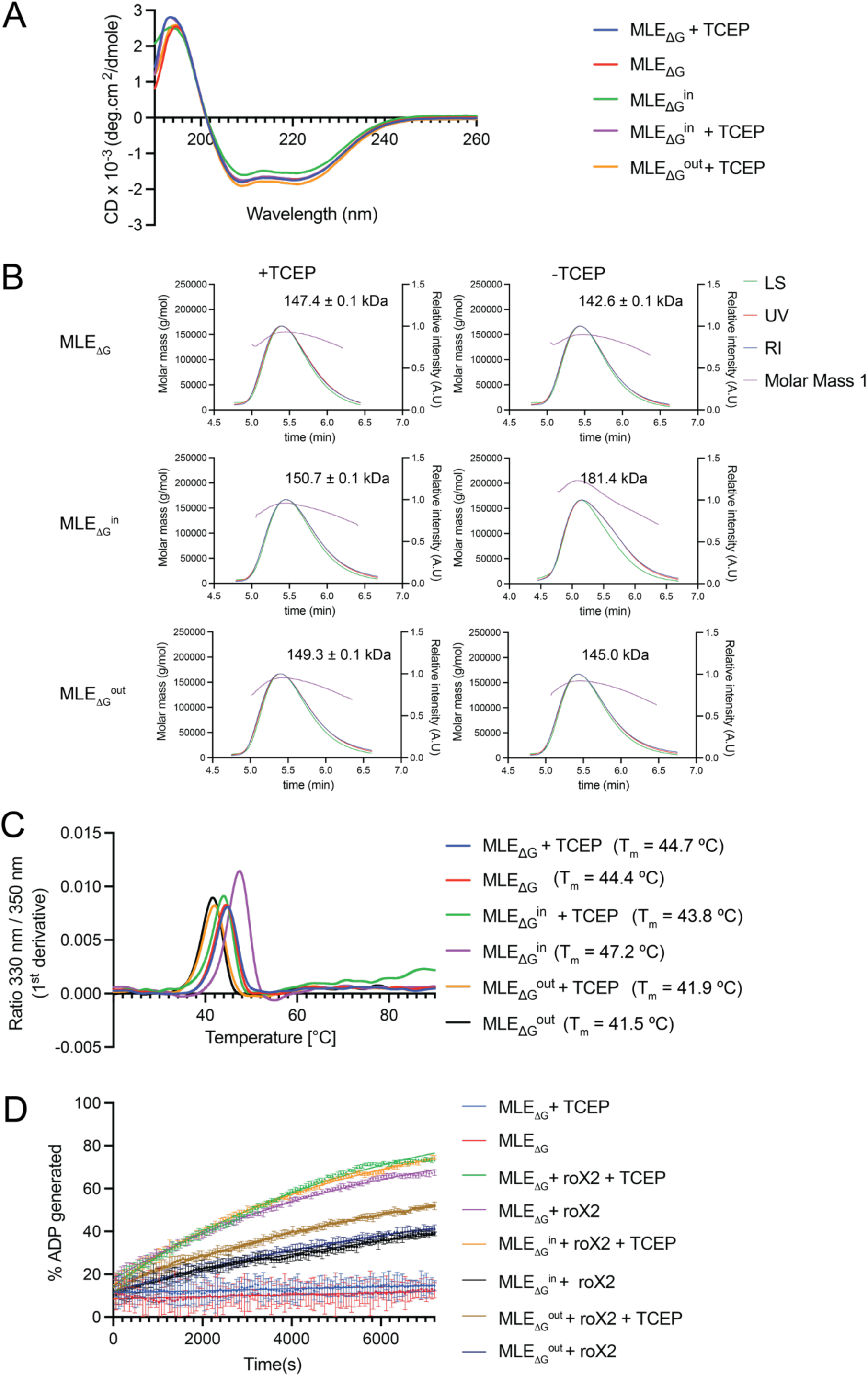
Biophysical characterization of MLE mutants. (A) Circular dichroism spectroscopy curves for MLE_ΔG_ and mutants revealing no prominent changes in secondary structure of the mutants compared to the wild-type protein. (B) SEC-MALS curves for MLE_ΔG_ and its mutants in the presence and absence of TCEP along with the experimentally determined molecular weights. The expected molecular weights for MLE_ΔG_ and MLE_ΔG_^in^ is 129.7 kDa and for MLE_ΔG_^out^ is 128.9 kDa. (C) Nano-DSF curves for MLE and its mutants along with the melting temperature of the proteins are presented. MLE_ΔG_^in^ shows a slight increase in melting temperature in the absence of TCEP because of the stabilizing effect of the cysteine bond formation on the MLE structure. (D) NMR ATPase assays of MLE_ΔG_ and mutants in absence or presence of roX2 RNA and TCEP, respectively. The percentage of ADP generated over time is plotted.

**Figure S7:**
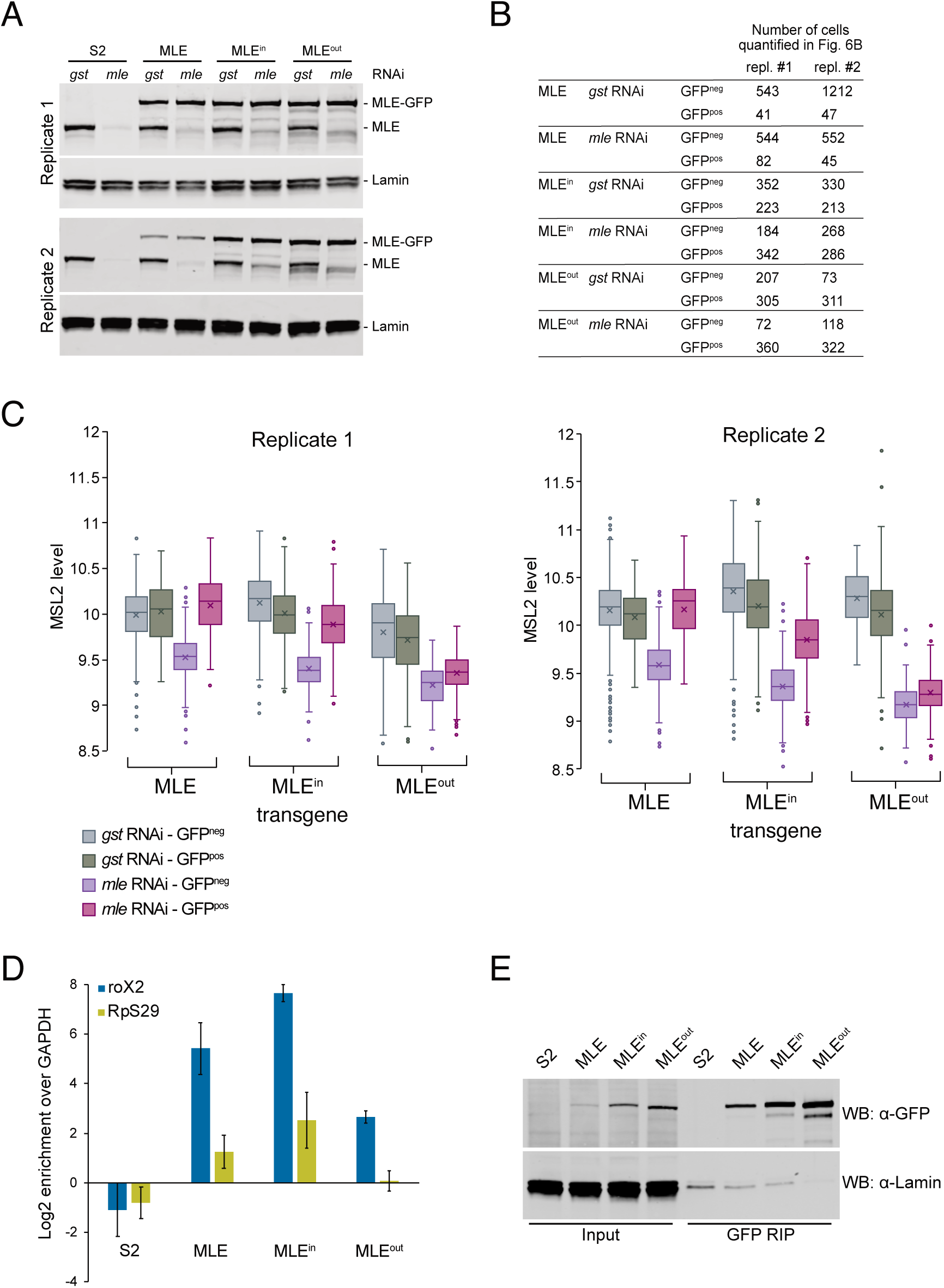
Restricting dsRBD2 conformation affects X chromosome localisation, territory formation and RNA binding *in vivo*. (A) RNAi efficiency in S2 cells stably expressing RNAi-resistant MLE_fl_-eGFP wild type and mutants used for immunostaining experiments in Figure 6. Cells were treated with control (*gst)* dsRNA or with dsRNA targeting endogenous *mle* and were analyzed in Western blot using an MLE-specific antibody, which detects endogenous and transgenic MLE_fl_-eGFP. Non-transfected S2 (‘S2’) cells were treated the same way. Lamin served as loading control. Both independent replicates analyzed in Figure 6 are presented. (B) Number of cells per replicate, which do (GFP^pos^) or do not (GFP^neg^) express MLE_fl_-eGFP transgenes and were included in the quantification shown in Figure 6B. (C) Boxplot of the immunofluorescence-based complementation assay. Shown are the nuclear MSL2 levels in GFP-positive (expressed MLE_fl_-eGFP wild type or mutant transgene) and GFP-negative cells after *gst* or *mle* RNAi treatment. The number of cells analyzed is given in S7B. The data of two independent replicates are presented. (D) RNA immunoprecipitation (RIP) of GFP-tagged MLE_fl_-wild type and mutants from stable S2 cells. Relative enrichment (IP/input) of roX2 and RpS29 transcripts was analyzed by RT-qPCR and is presented normalized to unbound GAPDH. Error bars represent the standard deviation for four independent replicates. (E) Western blot analysis of the RNA immunoprecipitation efficiency of MLE_fl_-eGFP wild type and mutants stably expressed in S2 cells. ‘S2’ represents a GFP-RIP control using non-transfected S2 cell extract. Protein levels in input and GFP-immunoprecipitated fractions were detected using anti-GFP antibody. Lamin served as loading control. One representative example out of four replicates is shown.

