## Supplementary Tables for "Structural basis of RNA-induced autoregulation of the DExH-type RNA helicase maleless"

**Key resources table**

| REAGENT or RESOURCE | SOURCE | IDENTIFIER |
| --- | --- | --- |
| Antibodies | | |
| Mouse anti-GFP | Roche | 11814460001 |
| Rat anti-MLE 6E11 | (Izzo et al., 2008) | NA |
| Rabbit anti-MSL2 | (Straub et al., 2005) | NA |
| Mouse anti-Lamin (T40) | H. Saumweber | NA |
| Donkey anti-mouse-Alexa488 | Jackson Immuno Research | 715-545-151 |
| Donkey anti-rat-Cy3 | Jackson Immuno Research | 712-165-153 |
| Donkey anti-rabbit-Alexa647 | Jackson Immuno Research | 711-605-152 |
| ChromoTek GFP-Trap Agarose | proteintech | gta |
| Bacterial and virus strains | | |
| E. coli chemically competent BL21(DE3) | ThermoFisher | C600003 |
| E. coli chemically competent Dh5a-T1R | ThermoFisher | 12297016 |
| E. coli electrocompetent DH10Bac | ThermoFisher | 10361012 |
| Biological samples |  |  |
| Chemicals, peptides, and recombinant proteins | | |
| Adenylyl-imidodiphosphate AMP-PNP | Roche | 10102547001 |
| Bis(sulfosuccinimidyl)suberate BS3 crosslinker | Thermo Fisher | A39266 |
| pCp-Cy5 | Jena Bioscience | NU-1706-Cy5 |
| Sf-900™ II SFM | ThermoFischer scientific | 10902088 |
| T4-RNA ligase | ThermoFischer scientific | EL0021 |
| Gibco Schneider’s Drosophila medium | ThermoFischer scientific | 21720001 |
| Fetal calf serum | Sigma-Aldrich | F7524 |
| SDB-RPS stage tip material | 3 M Empore | PN 2241 |
| Trypsin/Lys-C Mix, Mass Spec Grade | Promega | V5073 |
| Critical commercial assays | | |
| Deposited data | | |
| Cryo EM structure of MLE_ΔG_^apo^ | This study | PDB ID: 8B9L  EMDB ID: 15935 |
| Cryo EM structure of MLE_ΔG_ | This study | PDB ID: 8B9J  EMDB ID: 15933 |
| Cryo EM structure of MLE_ΔG_+SL7 RNA | This study | PDB ID: 8B9K  EMDB ID: 15934 |
| Cryo EM structure of MLE_ΔG_+U10 RNA | This study | PDB ID: 8B9G  EMDB ID: 15931 |
| Cryo EM structure of MLE_ΔG_+UUC RNA | This study | PDB ID: 8B9I  EMDB ID: 15932 |
| Experimental models: Cell lines | | |
| Drosophila melanogaster S2 cells, subclone L2-4 | Patrick Heun, Edinburgh |  |
| Experimental models: Organisms/strains | | |
| Oligonucleotides | | |
| MLE_257_NCOI_FP:TGAGCCATGGCGGATGAGCAGCTGAAGCCATATCCG | Sigma-Aldrich | NA |
| dsRBD2_145_NCOI_FP:TGAGCCATGGCGGAGCAGAGGGACATGAACGAAGCGGAG | Sigma-Aldrich | NA |
| MLE_FP_NCOI:TGAGCCATGGCGATGGATATAAAATCTTTTTTGTACCAATTTTGTG | Sigma-Aldrich | NA |
| E195C_FP:ATATACACCAGTGGGCCCGTGCCACGCTAGGAGCTTTTTGG | Sigma-Aldrich | NA |
| E195C_RP:CCAAAAAGCTCCTAGCGTGGCACGGGCCCACTGGTGTATAT | Sigma-Aldrich | NA |
| S633C_FP:TGGCCATGCTATCCGAATGCGACGTTAGCTTTGAGC | Sigma-Aldrich | NA |
| S633C_RP:GCTCAAAGCTAACGTCGCATTCGGATAGCATGGCCA | Sigma-Aldrich | NA |
| R590E602D603E605E608_AAAAA_FP: AGTCGCGCCGAAAGGCCAAGGAAGTGGAGGACGAGGAGCAATTGCTTTCCGCGGCCAAGGCCGAGGCGGCAATCAACTATAACAAGGTGTG | Sigma-Aldrich | NA |
| CACACCTTGTTATAGTTGATTGCCGCCTCGGCCTTGGCCGCGGAAAGCAATTGCTCCTCGTCCTCCACTTCCTTGGCCTTTCGGCGCGACT | Sigma-Aldrich | NA |
| D634A_FP:ATGCTATCCGAATCGGCCGTTAGCTTTGAGCTG | Sigma-Aldrich | NA |
| D634A_RP:CAGCTCAAAGCTAACGGCCGATTCGGATAGCAT | Sigma-Aldrich | NA |
| H746N747T754_AAA_FP:GCGCATGAAGCTCTTTACTTCAGCTGCCAACCTAACCAGCTACGCCGCAGTTTGGGCAAGCAAAACCAATTTGG | Sigma-Aldrich | NA |
| H746N747T754_AAA_RP:CCAAATTGGTTTTGCTTGCCCAAACTGCGGCGTAGCTGGTTAGGTTGGCAGCTGAAGTAAAGAGCTTCATGCGC | Sigma-Aldrich | NA |
| E790A_FP:GCCCGCTTCCAAGCGCTAGCGGACAATCTTACGCCGGAGATG | Sigma-Aldrich | NA |
| E790A_RP:CATCTCCGGCGTAAGATTGTCCGCTAGCGCTTGGAAGCGGGC | Sigma-Aldrich | NA |
| E835A_FP:CCTCCGGTAGACGCAGTAATCGCAGCTGAGGTGTTGCTTCGCGAG | Sigma-Aldrich | NA |
| E835A_RP:CTCGCGAAGCAACACCTCAGCTGCGATTACTGCGTCTACCGGAGG | Sigma-Aldrich | NA |
| K1027A_FP:GCGCAAAGTGCTGACTACAGAGTCTGCAGCAGCGTTACTGCACAAAACCTCGG | Sigma-Aldrich | NA |
| K1027A_RP:CCGAGGTTTTGTGCAGTAACGCTGCTGCAGACTCTGTAGTCAGCACTTTGCGC | Sigma-Aldrich | NA |
| R1057A_FP:CTTCGTTTTCGGCGAGAAGATTGCCACGCGAGCTGTTTCCTGCAAG | Sigma-Aldrich | NA |
| R1057A_RP:CTTGCAGGAAACAGCTCGCGTGGCAATCTTCTCGCCGAAAACGAAG | Sigma-Aldrich | NA |
| RNA sequence: SL7mod-UUC: GUGUAAAAUGUUGCU AGCAAAUAUAUAUGCUAGUAACGUUUUACGCCCUCUUUCUUUCUU | Integrated DNA Technologies | NA |
| RNA sequence: SL7-up-BHQ-1:  GUGUAAAAUGUUGCUAGCA-BHQ1 | Biomers | NA |
| RNA sequence: SL7-down-6FAM: 6-FAM-UGCUAGUA ACGUUUUACGCCCUCUUUCUUUCUU | Biomers | NA |
| RNA sequence: SL7-up: GUGUAAAAUGUUGCUAGCA | Biomers | NA |
| RNA sequence: U10 RNA: UUUUUUUUUU | Biomers | NA |
| RNA sequence: UUC RNA:CCUCUUUCUUUC | Biomers | NA |
| roX2-SL7.fw:  GACGTGTAAAATGTTGCAAATTAAG | Biomers | (Maenner et al., 2013) |
| roX2-SL7.rv:  TGACTGGTTAAGGCGCGTA | Biomers | (Maenner *et al.*, 2013) |
| RpS29.fw:  AGCGCATCGAAGCATTGATT | Biomers | This study |
| RpS29.rv:  GCGGAAGGTGGTAACTGTTG | Biomers | This study |
| 7SK.fw:  GATAACCCGTCGTCATCCAG | Biomers | (Hallacli et al., 2012) |
| 7SK.rv:  AGTAATTCTGCCTGGCGTTG | Biomers | (Hallacli *et al.*, 2012) |
| GAPDH.fw:  GGAGCCACCTATGACGAAAT | Biomers | (Quinn et al., 2014) |
| GAPDH.rv:  GTAGCCCAGGATTCCCTTC | Biomers | (Quinn *et al.*, 2014) |
| *mle* RNAi.fw:  TTAATACGACTCACTATAGGGAGAATGGATATAAAATCTTTTTTGTACCAATTTTG | Sigma-Aldrich | (Straub et al., 2008) |
| *mle* RNAi.rv:  TTAATACGACTCACTATAGGGAGAACAGGGCGCATGACTTGCT | Sigma-Aldrich | (Straub *et al.*, 2008) |
| *gst* RNAi.fw:  TTAATACGACTCACTATAGGGAGAATGTCCCCTATACTAGGTTA | Sigma-Aldrich | (Straub *et al.*, 2008) |
| *gst* RNAi.rv:  TTAATACGACTCACTATAGGGAGAACGCATCCAGGCACATTG | Sigma-Aldrich | (Straub *et al.*, 2008) |
| Recombinant DNA | | |
| pFastBac-MLE-flag | This study |  |
| pFastBac-His-MLE_ΔG_ | This study |  |
| pFastBac-His-MLE (257-1158) | This study |  |
| pFastBac-MLE^in^-flag | This study |  |
| pFastBac-MLE^out^-flag | This study |  |
| pFastBac-His-MLE_ΔG_^in^ | This study |  |
| pFastBac-His-MLE_ΔG_^out^ | This study |  |
| pFastBac-His-MLE_ΔG_ | This study |  |
| pHsp70-MLE-eGFP | This study |  |
| pHsp70-MLE^in^-eGFP | This study |  |
| pHsp70-MLE^out^-eGFP | This study |  |
| pET-M11-Thioredoxin-dsRBD1,2 | (Jagtap et al., 2019) |  |
| Software and algorithms | | |
| Phenix | (Adams et al., 2010) | SBGrid Consortium |
| Coot | (Emsley and Cowtan, 2004) | SBGrid Consortium |
| Crossfinder | (Forne et al., 2012; Mueller-Planitz, 2015) |  |
| Cryosparc | (Punjani et al., 2017) | https://cryosparc.com/ |
| Relion | (Scheres, 2012) | https://github.com/3dem/relion |
| Pymol | Schrodinger, LLC | https://pymol.org/2/ |
| Proteome discoverer 2.2 | Thermo scientific |  |
| GraphPad Prism 9 | GraphPad software Inc. | https://www.graphpad.com/scientificsoftware/prism/ |
| CcpNMR | (Skinner et al., 2016; Vranken et al., 2005) | https://ccpn.ac.uk/software/version-2/ |
| Adobe Illustrator | Adobe | https://www.adobe.com/products/illustrator.html |
| NMRPipe | (Delaglio et al., 1995) | SBGrid Consortium |
| Fiji | (Schindelin et al., 2012) |  |
| xvis web | (Grimm et al., 2015) | <https://xvis.genzentrum.lmu.de> |
| Other | | |
| 25 cm x 75 µm ID, 1.6 µm C18 | IonOpticks | AUR2-25075C18A-CSI |
| ReproSil-Pur C18-AQ 2.4 μm | Dr. Maisch | r124.aq. |
| RSLCnano Ultimate 3000 system | Thermo Fisher | ULTIM3000RSLCNANO |

Table 1: Cryo-EM data collection, refinement, and validation statistics

|  | MLE_ΔG_^apo^ | MLE_ΔG_ | MLE_ΔG_+ U10 | MLE_ΔG_+UUC | MLE_ΔG_+SL7  dts1 | MLE_ΔG_+SL7  dts2 |
| --- | --- | --- | --- | --- | --- | --- |
| EMDB ID | 15935 | 15933 | 15931 | 15932 | 15934 | |
| PDB ID | 8B9L | 8B9J | 8B9G | 8B9I | 8B9K | |
| **Data collection and processing** | | | | | | |
| Magnification | 105,000 | 105,000 | 105,000 | 105,000 | 105,000 | 105,000 |
| Voltage (kV) | 300 | 300 | 300 | 300 | 300 | 300 |
| Electron exposure (e–/Å^2^) | 50.5 | 51.7 | 61.2 | 48.5 | 47.9 | 51.7 |
| Defocus range (μm) | -0.8 to -2.0 | -0.8 to -2.0 | -0.8 to -2.0 | -0.8 to -2.0 | -0.8 to -2.0 | -0.8 to -2.0 |
| Pixel size (Å) | 0.822 | 0.822 | 0.822 | 0.822 | 0.822 | 0.822 |
| Initial particle images (no.) | 1470793 | 3199267 | 4503260 | 1807333 | 1763219 | 184188 |
| Final particle images (no.) | 270599 | 234365 | 559766 | 476446 | 325471 | |
| Map resolution (Å) | 3.45 | 3.45 | 2.95 | 2.86 | 4.04 | |
| FSC threshold | 0.143 | 0.143 | 0.143 | 0.143 | 0.143 | |
| Map resolution range (Å) | 2.9-6.8 | 2.9-6.8 | 2.5-5.9 | 1.8-3.0 | 3.5-8.6 | |
| **Refinement** | | | | | | |
| Initial model used (PDB code) | 5AOR | MLE_ΔC_ | 5AOR | 5AOR | 5AOR, 6I3R | |
| Model resolution (Å) | 3.45 | 3.45 | 2.95 | 2.86 | 4.04 | |
| Model resolution range (Å) | 420.8-3.45 | 420.8-3.45 | 420.8-2.95 | 420.8-2.86 | 526.0-4.04 | |
| Map sharpening *B* factor (Å^2^) | -154 | -137 | -128 | -134 | -203 | |
| Model composition  Non-hydrogen atoms  Protein residues  Nucleotide residues  Ligands | 6827  861  0  0 | 6879  864  0  3 | 7801  953  10  3 | 8036  984  11  3 | 8522  966  40  2 | |
| *B* factors (Å^2^)  Protein  Ligand  Nucleotide | 45.3  --  -- | 45.3  43.2  -- | 40.5  21.6  52.1 | 41.1  30.1  45.57 | 92.1  20  53.0 | |
| R.m.s. deviations  Bond lengths (Å)  Bond angles (°) | 0.003  0.69 | 0.004  0.79 | 0.004  0.628 | 0.004  0.68 | 0.003  0.6 | |
| Validation  MolProbity score  Clashscore  Poor rotamers (%) | 2.2  14.3  0.26 | 2.12  13.31  0.39 | 1.68  9.02  1.31 | 2.11  11.72  2.31 | 2.81  16.79  4.71 | |
| Ramachandran plot  Favored (%)  Allowed (%)  Disallowed (%) | 90.74  8.44  0.82 | 91.94  7.59  0.47 | 97.46  2.43  0.11 | 96.32  3.58  0.10 | 89.09  10.4  0.52 | |

Table S1: Average intensity of dsRBD1, linker and dsRBD2 upon titration with helicase module and substrates. The errors represent standard deviation of the averaged intensity ratios.

|  | dsRBD1,2 + helicase module | dsRBD1,2 + helicase module + UUC RNA | dsRBD1,2 + helicase module + UUC RNA + ADP:AlF_4_ |
| --- | --- | --- | --- |
| dsRBD1 | 0.31 ± 0.2 | 0.32 ± 0.2 | 0.29 ± 0.2 |
| L1 linker | 1.04 ± 0.06 | 1.04 ± 0.06 | 0.80 ± 0.1 |
| dsRBD2 | 0.93 ± 0.1 | 0.94 ± 0.1 | 0.91 ± 0.2 |

Table S2: CLMS data for MLE apo and in complex with U10 and SL7 RNA. Crosslinks between lysine residues in MLE as identified by mass spectrometry. Position of lysine residues is given relative to the Start-Methionine. Empty cells indicate absence of crosslinks of the corresponding lysine residue. Intermolecular crosslinks of MLE can neither be excluded nor distinguished.

Table S3: IC_50_ of MLE and its mutants for dsRNA (roX2 SL678) and ssRNA (UUC RNA) as determined from FP assays. Errors represent standard error of fitting.

|  | roX2 SL678 (nM) | UUC RNA (nM) |
| --- | --- | --- |
| MLE_ΔG_ | 4 ± 0.6 | 302 ± 43 |
| MLE_ΔG_^in^ | 80 ± 11 | 280 ± 19 |
| MLE_ΔG_^out^ | 8.9 ± 1.9 | > 2500 |

Table S4: Rate constant for the rate of ADP produced during the ATPase activity of MLE_ΔG_ and its mutants as determined from NMR ATPase assays.

|  | +TCEP (s^-1^) | -TCEP (s^-1^) |
| --- | --- | --- |
| MLE_ΔG_ | 6.7 e^-5^ | 7.4 e^-5^ |
| MLE_ΔG_ + roX2 | 7.3 e^-6^ | 4.7 e^-6^ |
| MLE_ΔG_^in^ + roX2 | 3.9 e^-6^ | 1.0 e^-5^ |
| MLE_ΔG_^out^ + roX2 | 6.0 e^-6^ | 4.7 e^-6^ |

Table S5: Rate constant and the plateau values for the helicase activity of MLE_ΔG_ and its mutants determined from real time fluorescence RNA helicase assay.

|  | Plateau (%) | K_Fast_ (s^-1^) | K_Slow_ (s^-1^) |
| --- | --- | --- | --- |
| MLE_ΔG_ + TCEP | 46.15 | 0.02715 | 0.001142 |
| MLE _ΔG_ | 56.54 | 0.07122 | 0.001398 |
| MLE _ΔG_^in^ + TCEP | 36.78 | 0.0087 | 0.000529 |
| MLE _ΔG_^in^ | 18.28 | 0.01558 | 0.000507 |
| MLE _ΔG_^out^ + TCEP | 2.49 | Not defined | 0.000978 |
| MLE _ΔG_^out^ | 2.2 | Not defined | 0.001235 |
